# Anisotropic expansion of hepatocyte lumina enforced by apical bulkheads

**DOI:** 10.1101/2020.12.31.424978

**Authors:** Lenka Belicova, Urska Repnik, Julien Delpierre, Elzbieta Gralinska, Sarah Seifert, José Ignacio Valenzuela, Hernán Andrés Morales-Navarrete, Christian Franke, Helin Räägel, Evgeniya Shcherbinina, Tatiana Prikazchikova, Victor Koteliansky, Martin Vingron, Yannis Kalaidzidis, Timofei Zatsepin, Marino Zerial

## Abstract

Lumen morphogenesis is key to the function of organs and results from the integration of molecular pathways and mechanical forces^1–3^. The mechanisms governing anisotropic lumen expansion remain elusive^4–6^. In contrast to epithelial cells which have simple apico-basal polarity and form tubes, hepatocytes are multi-polar and form narrow lumina that grow anisotropically between adjacent cells, collectively generating a complex 3D network of bile canaliculi (BC)^7,8^. Here, we studied lumen elongation and BC morphogenesis in differentiating primary mouse hepatoblasts *in vitro*. Remarkably, we discovered a pattern of specific extensions of the apical membrane traversing the lumen between adjacent hepatocytes and sealed by tight junctions, reminiscent of the bulkheads of boats. These structures were also present in the developing liver. A targeted screen revealed that silencing of Rab35 caused loss of the bulkheads, conversion of hepatocyte into simple epithelial polarity and formation of spherical lumina *in vitro*. Strikingly, we could re-engineer hepatocyte polarity and tissue morphogenesis *in vivo* in the embryonic liver, converting BC into simple epithelial tubes. Our results suggest that the apical bulkheads of hepatocytes are cell-intrinsic anisotropic mechanical elements that ensure stability of the elongating lumen between two cells, thus determining the structure of BC during liver tissue morphogenesis.

## Main

*Lumen* morphogenesis is essential for several organs. *Lumina* are generated by epithelial cells that exhibit apico-basal polarity, with the apical surface facing the internal *lumen* and the basal surface contacting the basement membrane^9,10^. *Lumina* expand either isotropically yielding spherical structures (acini and alveoli *in vivo*, cysts and organoids *in vitro*) or anisotropically, generating a variety of epithelial tube shapes across tissues (e.g. lungs, intestine, kidney, liver). The anisotropic expansion of lumina is more difficult to explain than the isotropic one because it results from specific combinations of molecular pathways and physical forces^1,2,11^. Physical forces can act on the tissue or cellular level. The liver provides a good example for a variety of lumen morphogenesis that is essential for its function^12–16^. It contains two types of epithelial cells, bile duct cells (cholangiocytes) and hepatocytes, both derived from embryonic progenitors called hepatoblasts^13^. Bile duct cells have the typical apico-basal polarity and expand isotropically to form 3D cysts *in vitro*^17,18^, but generate tubes *in vivo*^19,20^, which elongate under tissue-level forces^2^. In contrast, hepatocytes are multi-polar and their apical surfaces elongate anisotropically as a tubular belt surrounding the cells. Hepatocytes can initiate apical *lumina* with multiple neighbouring hepatocytes concurrently in all directions, allowing them to form a complex 3D luminal network of highly branched ^~^ 1 μm-thin bile canaliculi (BC)^8^. The morphology of BC implies that the anisotropy of lumen elongation is not imposed by tissue-level forces but rather by local forces at the cellular level.

The mechanisms underlying the anisotropy of lumen formation at the cellular scale remain elusive. Models based mainly on *in vitro* studies propose cell division as a key determinant of lumen elongation^15,21,22^. In the developing liver however, as hepatoblasts differentiate into hepatocytes, they gradually stop proliferating^23^. Yet, an almost fully connected BC network is generated^20^, arguing for additional mechanisms driving lumen elongation. From the physics of thin-shells^24,25^, lumen elongation by foetal hepatocytes requires mechanisms based either on the anisotropic structure of the apical actomyosin cortex or some other mechanical elements to enforce a tubular lumen. Here, we set to identify such mechanisms.

### Anisotropic lumen morphogenesis by hepatocytes

We established a culture of primary mouse hepatoblasts isolated from embryonic livers based on Dlk1 expression^26^ to differentiate them into hepatocytes (see expression of mature hepatocyte markers Extended data Fig.3a). The cells generated elongated and branched tubular lumina enriched in F-actin, and positive for the apical marker CD13 and the tight junction protein ZO-1 (Figure 1a, b). This system therefore recapitulates *de novo* formation of branched BC-lumina *in vitro* similar to the developing liver *in vivo*.

**Figure 1:**
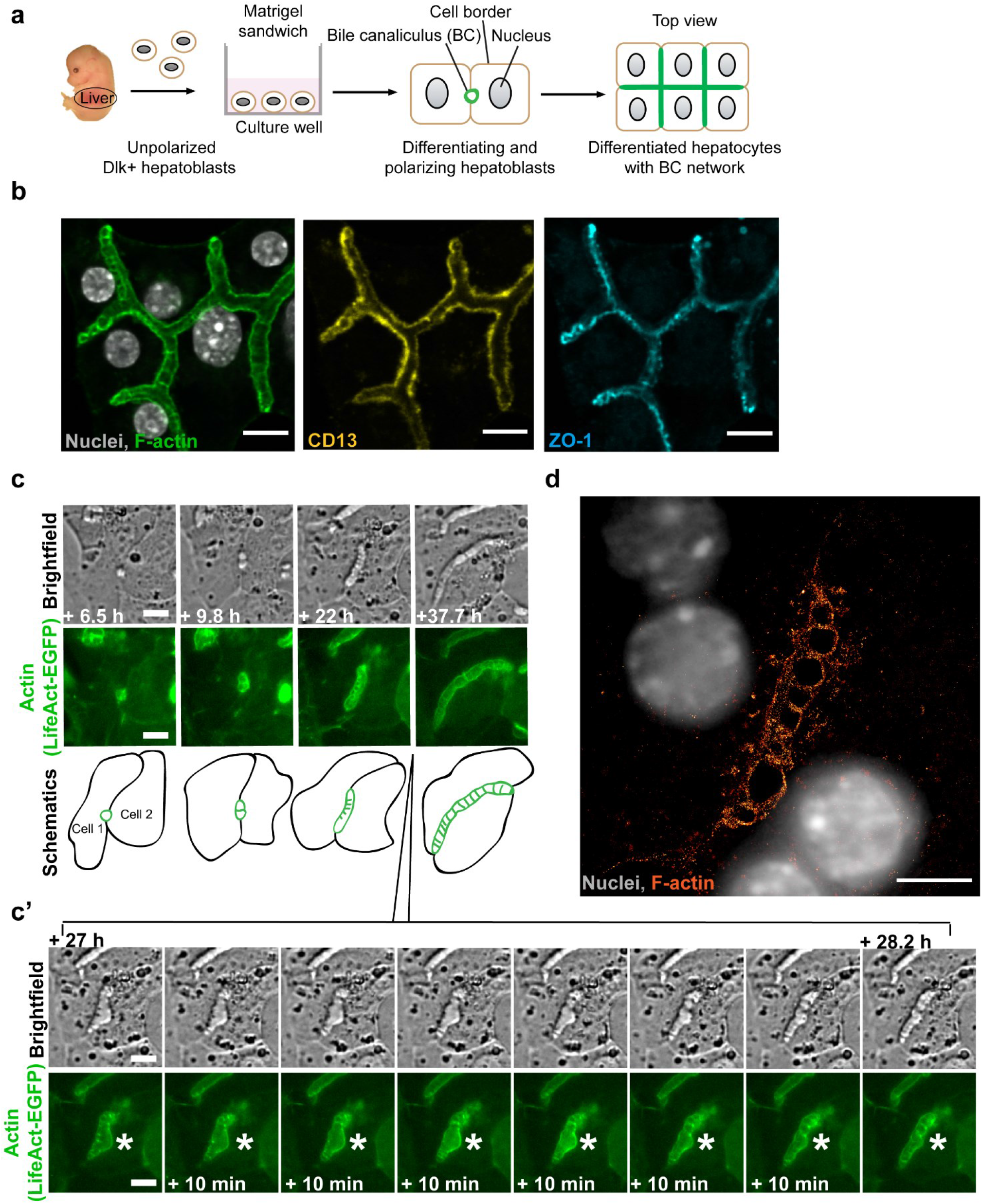
Lumen morphogenesis in hepatocytes is accompanied by specific actin structures that interconnect the two lumen-forming cells. **a**, Schematic overview of primary Dlk1+ hepatoblasts in Matrigel sandwich culture differentiating into hepatocytes and recapitulating BC formation. **b**, Differentiated hepatocytes form branched interconnected BC lumina. Immunofluorescence microscopy images of differentiated hepatocytes stained for F-actin with Phalloidin-Alexa488, and for apical markers CD13 and ZO-1. Scale bar: 10 μm. **c**, Images from live-cell time-lapse microscopy documenting the formation of BC between two differentiating hepatoblasts expressing LifeAct-EGFP. During imaging, the extending tubular lumen displayed a bulge at 27h from the start of imaging, which was subsequently “re-absorbed”. The insert **c’** shows individual frames taken each 10 min from this timepoint, documenting the recovery of the tubule (white star). Note the transverse striped pattern in brightfield and actin channels, which is apparent when the lumen is tubular but not observed within the bulge. Scale bar: 10 μm. See also Video 1. **e**, Single-molecule localization microscopy (SMLM) image of a lumen between two differentiated hepatocytes, actin labelled with Phalloidin-Alexa647. Note the transverse striped actin pattern. Scale bar: 5 μm.

To study lumen morphogenesis, we performed live-cell time-lapse microscopy on differentiating hepatoblasts stably expressing LifeAct-EGFP as actin label. We followed lumenogenesis for up to 52 hours and categorised four sequential steps: 1) lumen initiation, 2) elongation, 3) branching and 4) fusion. We frequently observed single cells initiating multiple individual lumina with their neighbours (Extended data Fig.1a, Video 2). After formation, lumina elongated into tubes until they spanned the entire cell–cell contact (Figure 1c, Extended data Fig. 1a-c, Videos 1-4). At this point, a lumen could fuse with another lumen (Extended data Fig.1b, left panels) or branch at a 3-cell contact (Extended data Fig.1b, right panels). The elongation of lumina occurred in the absence of cell division.

We were intrigued by the presence of dark stripes in the bright-field, transverse to the direction of lumen elongation (e.g. Figure 1c, Extended data Fig.1) which may correspond to high curvature of the apical membranes. The stripes also coincided with areas of high density of actin (LifeAct-EGFP). The pattern was evident early in lumen formation and continued as the lumina elongated, keeping a characteristic spacing between stripes (Figure 1c, Extended data Fig.1a-c). Interestingly, we also observed instances when the lumen transiently bulged outward, tending to a spherical lumen (Figure 1c (marked with a star), Video 1 and Extended data Fig.1c, Video 4). This coincided with the loss of the stripes. Subsequently, the tubular shape of the lumen recovered as new stripes formed, suggesting an active link between the striped pattern and lumen elongation.

To determine the micro-structure of the actin-rich stripes, we analysed the cortical F-actin labelled with Phalloidin-Alexa-647 using single-molecule localization microscopy (SMLM) on fixed, *in vitro* differentiated hepatocytes. Strikingly, we observed a *quasi*-periodic pattern of F-actin structures apparently crossing the lumen between two cells (Figure 1d), similar to the pattern of stripes in the bright-field (Figure 1c). Because the SMLM has a z-resolution of ^~^500 nm and these structures are >1 μm in height, we can conclude that the F-actin projects into the BC lumen and does not correspond to rings around it, e.g. as in the Drosophila tracheal tube^27,28^.

### Ultra-structural analysis reveals bulkhead-like apical transversal structures in the BC lumen

Given the presence of both actin filaments and tight junctions (ZO-1) traversing the lumen, we investigated these structures in greater detail by electron microscopy (EM) on serial sections and 3D reconstructions of the entire lumen volume. Remarkably, the EM section of Figure 2a shows a branched lumen between three hepatocytes, whose surfaces are connected by finger-like membrane processes. The fingers of one cell touch, or invaginate into, the opposing cell (Figure 2b) and the contact surfaces are sealed by tight junctions (Figure 2c, d). Interestingly, we often observed vesicles accumulated at the base of these processes.

**Figure 2:**
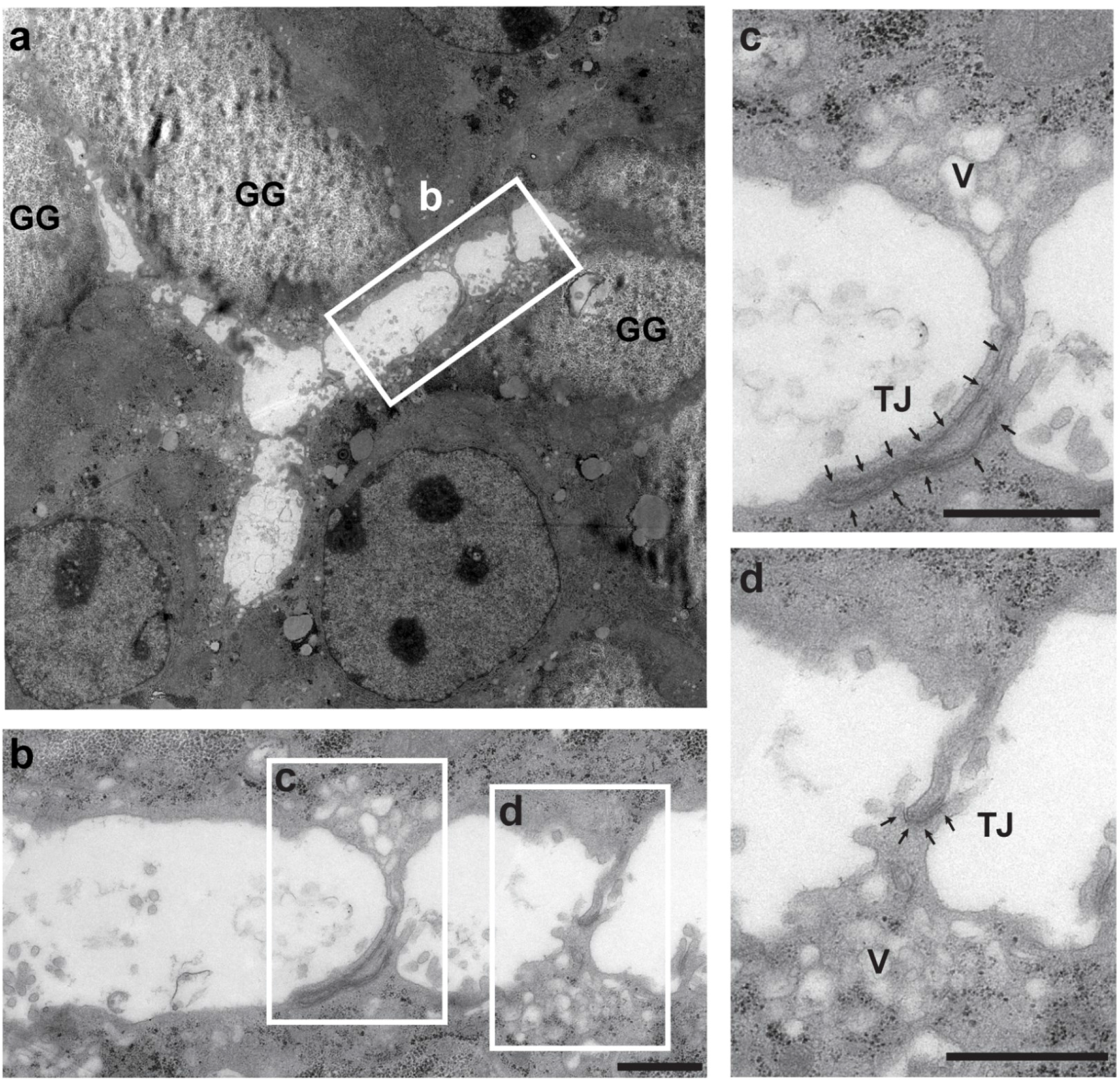
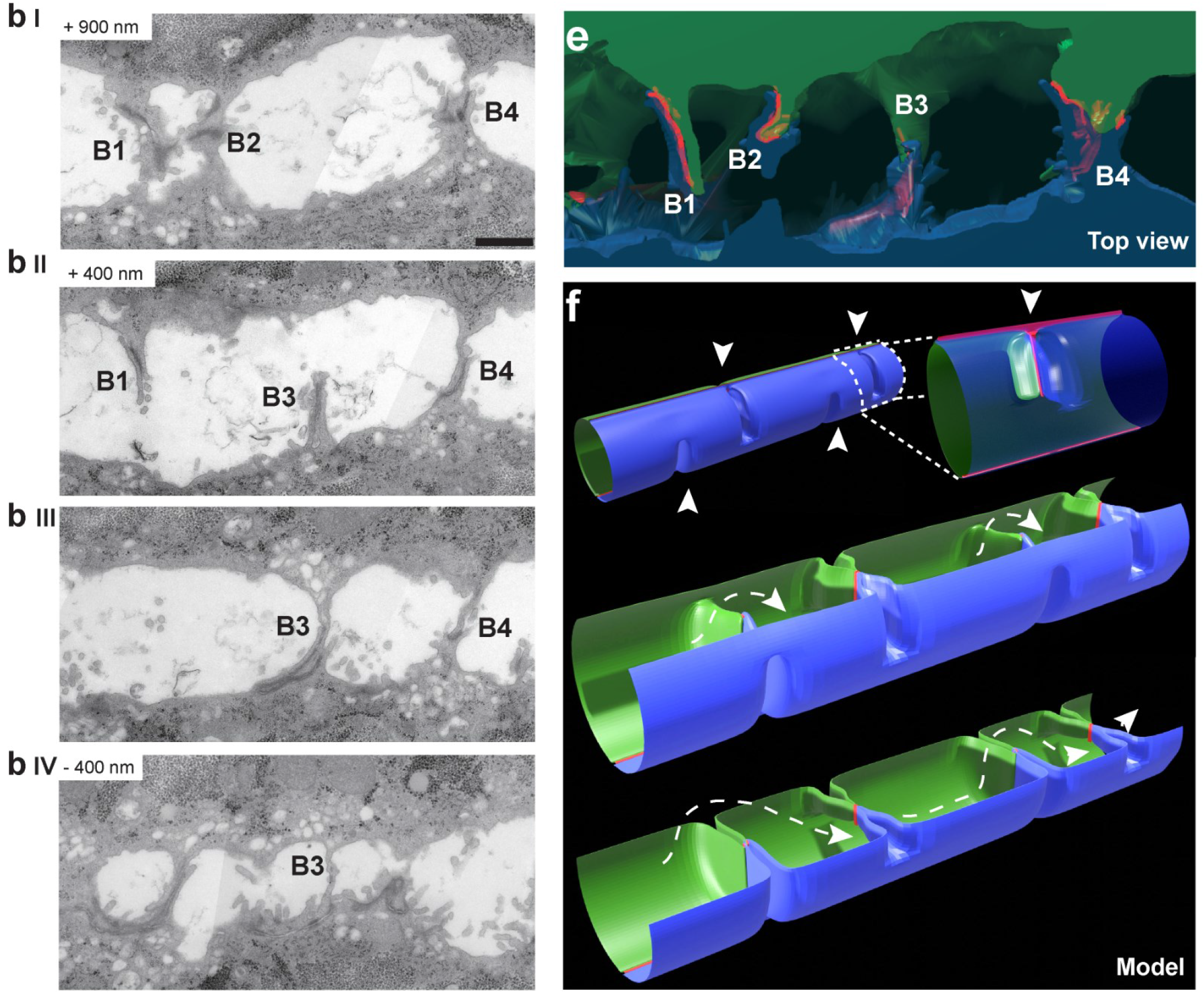
Ultra-structural analysis reveals a bulkhead-like pattern of transversal structures sealed by T-shaped tight junctions in the BC lumen. **a**, Electron microscopy image of a BC branched between three cells. Longitudinal section of *in vitro* differentiated hepatocytes. GG: glycogen granules. The outlined region is shown in panel **b**. Scale bar: 5 μm. **b**, Section of BC formed by two cells from a series of longitudinal 90-nm sections shown in b I-IV. The highlighted regions with transversal membrane connections are shown in the panel **c** and **d**. The membrane connections are not visible in every section **(b I-IV)**. Scale bar: 1 μm. **c**, and **d**, Detailed view of the outlined regions in panel **b**. Membrane connections are formed by apical surfaces of both cells lining the BC and include the tight junctions (TJ, arrows). Vesicles (V) are often observed to accumulate in the vicinity of the connections. Scale bar: 1 μm. **e**, 3D reconstruction of serial sections in **b (b I-IV)** based on apical plasma membranes and tight junctions rendering. The cytoplasm of the lumen-forming cells is in green and blue, the tight junctions are highlighted in red. See also Video 5. **f**, Simplified model of BC based on the 3D reconstruction in **e**with periodic bulkhead-like membrane connections formed from the top or the bottom of the lumen (arrowheads). The tight junctions (red) have a T-shape, with the junctions longitudinal along the tube connected with the junctions extending along the ridgeline within each bulkhead. Uninterrupted flow within the lumen between bulkheads is shown with a dotted line. See also Video 6.

From a single section it is impossible to establish whether the lumen is continuous or divided into separate chambers. The 3D reconstruction (Figure 2e, Video 5) revealed that the transversal finger-like processes (Figure 2a) were sections of structures resembling the bulkheads of a boat. The bulkheads consisted of two parts, each contributed by the apical surface of one of the two adjacent cells which formed a ridge-shaped process (see 3D model Figure 2f, Video 5 and 6). Importantly, the two ridges were sealed by tight junctions which followed an unusual T-shape, with the horizontal bar representing the junctions longitudinal along the tube and the vertical bar the junctions extending along the ridgeline (see scheme in Figure 2e, Video 5 and 6). The EM data are consistent with the presence of ZO-1 structures in the stripes crossing the lumen (Figure 1b). In some cases, the opposing processes are not precisely aligned along the ridgeline but shifted, forming a wide tight junction contact belt (Figure 2b, 2e, bulkheads B1, 4). The bulkheads can come either from the bottom (see Figure 2f, bulkhead B3) or the top of the tube (Figure 2b I-IV, e, bulkheads B1, 2, 4), but never intersect it completely, thus ensuring lumen continuity in the BC (Video 5 and 6). Consequently, from the 3D reconstruction of Figure 2b and Video 5, one can appreciate that the lumen has a tortuous shape. This accounts for the impression that the F-actin fluorescent and bright-field stripes only partially cross the lumen (Figure 1c). The bulkheads showed a *quasi*-periodicity similar to the pattern in the bright-field and of actin observed by live-cell imaging (Figure 1c) and SMLM (Figure 1d).

To rule out that these structures are an artefact of the in vitro system, we examined the embryonic (E15.5) liver by EM. Also here we could confirm the presence of the repetitive pattern of bulkhead-like transversal connections in the nascent BC (Extended data Fig.2a-b). The lumen shape was even more complex than *in vitro*, due to the 3D organization of the tissue, with a higher degree of freedom for cell-cell contacts. Importantly, also *in vivo* the bulkheads did not divide the BC lumen into isolated chambers (Extended data Fig.2c-d).

### Conversion of hepatocyte polarity into simple apico-basal polarity

The bulkhead-like apical processes could be a specific feature of hepatocyte polarity to enable the anisotropy of apical lumen growth. If so, their loss may convert hepatocyte polarity into simple epithelial polarity and induce the formation of cysts. Our *in vitro* system enables both types of polarity simultaneously, side by side in the same culture. The hepatoblasts that differentiate into hepatocytes form branched BC-like structures at the bottom of the well (Extended data Fig.3b), whereas the bile duct (Sox9+, EpCAM+) cells form 3D cysts raising into the medium (Extended data Fig.3b-c).

Therefore, to identify genes required for hepatocyte polarity, we performed a focused siRNA screen on 25 candidate genes, encoding key regulatory components of cell polarity (Table 1): apical junction formation (e.g. Pard3, Tjp1, Cldn2), cytoskeleton regulation (e.g. Mark2/Par1b, Stk11/Lkb1, Cdc42) and polarized trafficking (e.g. Rab11a, Rab35, Cdc42), including genes previously associated with the regulation of hepatocyte polarity^21,29–31^. Hepatoblasts were transfected with the siRNAs and after 5 days in culture stained for F-actin, which is enriched at the apical domain (Figure 1b-c). Hit candidates were those yielding a penetrant lumen phenotype with a minimum of two siRNAs. Silencing of *Ocln* (Extended data Fig.4a) and *Tjp1* (Extended data Fig.4b) yielded a loss of cell polarity, with de-localized F-actin and the apical marker CD13 due to the absence of lumen. Remarkably, out of the 25 genes screened, downregulation of Cdc42 and Rab35 did not compromise cell polarity, as judged by the localization of CD13 and ZO-1, but altered lumen morphology (Figure 3a-b, Extended data Fig.4c-d). *Cdc42* silencing caused dilated spherical lumina (Extended data Fig.4c), however, Rab35 knock-down yielded the most striking phenotype, causing the appearance of simple epithelial tubes (white arrowhead Figure 3b) and large cyst-like structures (yellow arrowhead Figure 3b) instead of BC (as in Luciferase, siLuc) (Figure 3a-b, Extended data Fig.4d, and Video 7). Optical sectioning and 3D reconstruction showed that the lumina of these tubes and cysts were connected with the BC formed by the neighbouring hepatocytes (Figure 3b), indicating that the cells indeed displayed the two types of polarity side by side, as in the peri-portal zone of liver tissue ^13,14,19^.

**Table 1:** Genes included in the focused siRNA screen.

| Gene name | Gene ID | Lumen phenotype | Number of siRNAs with phenotype | Category |
| --- | --- | --- | --- | --- |
| *Tjp1* | 21872 | Loss of polarity | 4/6 | Apical junctions |
| *Ocln* | 18260 | Loss of polarity | 2/6 | Apical junctions |
| *Cldn2* | 12738 | None | 0/6 | Apical junctions |
| *Gjb1* | 14618 | None | 0/6 | Apical junctions |
| *Pard3* | 93742 | None | 0/6 | Apical junctions/ Cytoskeleton |
| *Mark2 (Par1b)* | 13728 | None | 0/6 | Cytoskeleton |
| *Kif13b* | 16554 | None | 0/6 | Cytoskeleton |
| *Stk11 (Lkb1)* | 20869 | None | 0/6 | Cytoskeleton |
| *Cdc42* | 12540 | Spherical lumina | 3/6 | Cytoskeleton/ Polarized trafficking |
| *Arf6* | 11845 | Cyst-like lumina | 1/6 | Cytoskeleton/ Polarized trafficking |
| *Rab35* | 77407 | Cyst-like lumina | 5/6 | Cytoskeleton/ Polarized trafficking |
| *Rab4a* | 19341 | None | 0/6 | Polarized trafficking |
| *Rab4b* | 19342 | None | 0/6 | Polarized trafficking |
| *Rab8a* | 17274 | None | 0/6 | Polarized trafficking |
| *Rab8b* | 235442 | None | 0/6 | Polarized trafficking |
| *Rab10* | 19325 | None | 0/6 | Polarized trafficking |
| *Rab11a* | 53869 | None | 0/6 | Polarized trafficking |
| *Rab11b* | 19326 | None | 0/6 | Polarized trafficking |
| *Rab13* | 68328 | None | 0/6 | Polarized trafficking |
| *Rab14* | 68365 | None | 0/6 | Polarized trafficking |
| *Rab17* | 19329 | None | 0/6 | Polarized trafficking |
| *Rab21* | 216344 | None | 0/6 | Polarized trafficking |
| *Rab27a* | 11891 | None | 0/6 | Polarized trafficking |
| *Rab27b* | 80718 | None | 0/6 | Polarized trafficking |
| *Mal2* | 105853 | None | 0/6 | Polarized trafficking |

**Figure 3:**
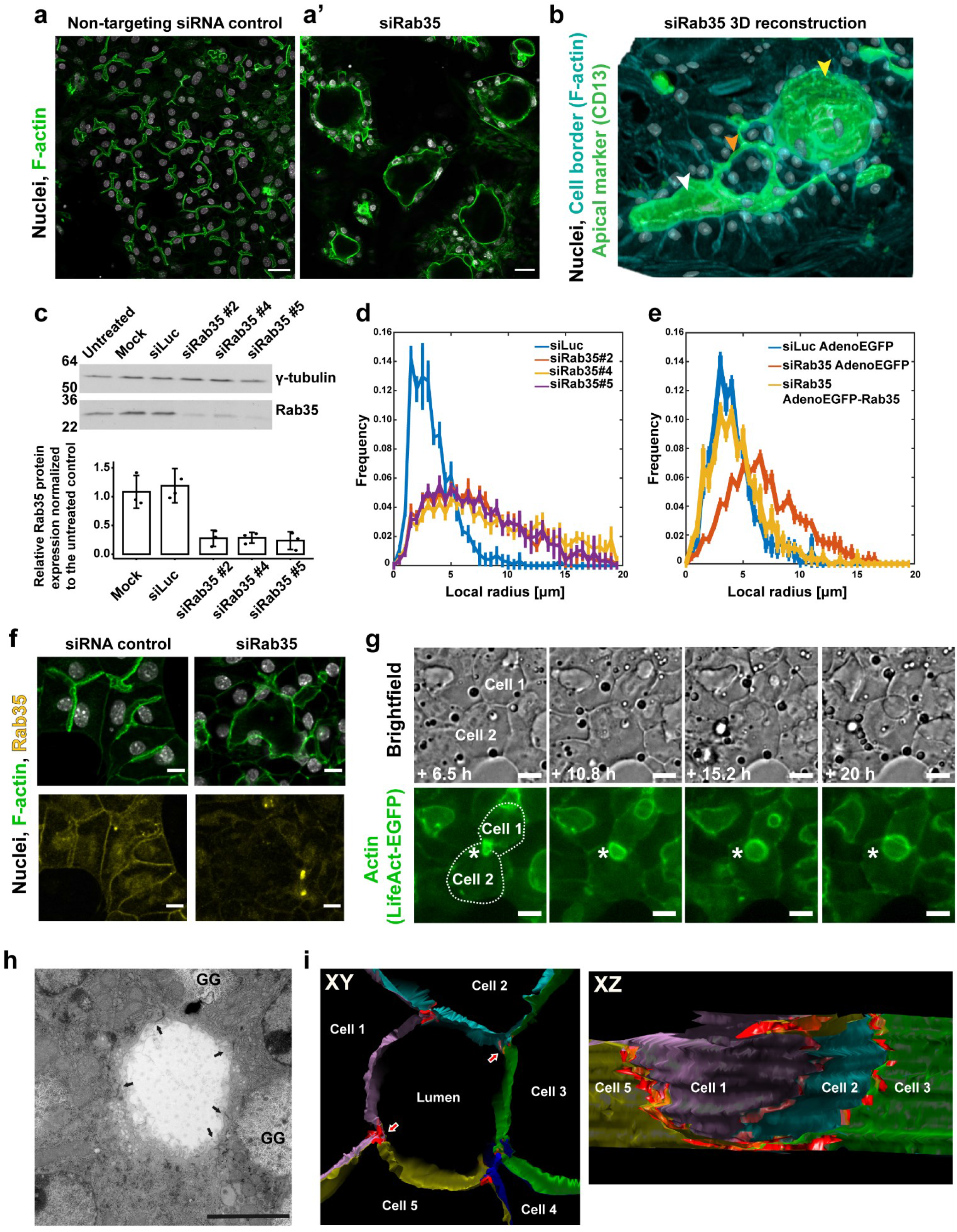
Silencing of Rab35 causes the loss of the transversal apical membrane bulkheads and formation of spherical cysts. **a**, Knock-down of Rab35 in differentiating hepatoblasts caused formation of cyst-like structures (**a’**), whereas cells treated with the control siRNA (siLuc) were unaffected and formed BC. Microscopy images of cells stained for F-actin with Phalloidin-Alexa488. Scale bar: 30 μm. See also Video 7. **b**, 3D reconstruction of the cells treated with Rab35 siRNA show the variability of the phenotype from enlarged swollen lumina of epithelial tubes (white arrowhead) to spherical cyst-like structures (yellow arrowhead) growing in the z-direction (lumina stained with the apical marker CD13). The BC (orange arrowhead) connected to the cyst, reminiscent of the organization of the peri-portal zone in liver tissue. **c**, Three independent Rab35 siRNA duplexes down-regulate Rab35 protein levels by 73% ± 3% (n = 3, error bars: SD). Representative Western blot and quantification of protein knock-down. **d**, Histogram of the local lumen radius in control cells and cells treated with Rab35 siRNA estimated based on microscopy image analysis. Rab35 knock-down by three independent siRNAs results in the shift towards the lumina with larger radius. Percentage of lumina > 6μm: siLuc 1.00±0.46%, siRab35#2: 20.55±1.66%, siRab35#4: 24.86±1.46%, siRab35#5: 19.95±2.81% (n= 3 (with 4 images per condition), error bars: SEM). **e**, The enlarged lumina phenotype in the cells treated with Rab35 siRNA is rescued by expression of human Rab35-EGFP from recombinant adenovirus. The frequency curve of the lumen radius (yellow) overlaps with the one of the control cells expressing EGFP only (blue). EGFP alone does not affect the lumen enlargement caused by Rab35 knock-down (red). Percentage of lumina > 6μm: siLuc AdenoEGFP: 2.39±0.44%, siRab35 AdenoEGFP: 22.35±1.08%, siRab35AdenoEGFP-Rab35: 3.33±1.05% (n= 3 (with 4 images per condition), error bars: SEM). **f**, Microscopy images of differentiated hepatocytes treated with Luciferase or Rab35 siRNA stained with Rab35 antibodies (yellow). Rab35 localizes to the apical and lateral plasma membrane and cytoplasmic puncta. The levels of Rab35 are markedly reduced in the cyst-like structures formed upon Rab35 siRNA transfection. Scale bar: 10 μm. **g**, In the cells treated with Rab35 siRNA, the lumina tend to grow as spheres instead of elongating as tubes. Images from the live-cell time-lapse microscopy experiment showing two neighbouring differentiating hepatoblasts expressing LifeAct-EGFP under Rab35 siRNA conditions. The white star indicates the forming lumen between the two cells. Note that the typical transverse striped actin pattern observed in the tubular BC is absent. Scale bar: 10 μm. See also Video 8. **h**, EM analysis of a cyst-like lumen resulting from the Rab35 knock-down. A longitudinal 90-nm section of the lumen formed between five cells. The bulkheads typical for hepatocyte BC are absent in the lumen. Arrows indicate tight junctions. GG: glycogen granules. Scale bar: 5 μm. **i**, Longitudinal view (left panel) through the middle of a 3D model of the lumen based on rendering plasma membranes and tight junctions (red) on serial sections. The five cells forming the lumen are represented in different colours. Red arrows point to the tight junctions at which the cyst is cut open to reveal the sagittal view (right panel). The lumen has a circular profile and tight junctions do not protrude into the lumen.

### Rab35 is rate-limiting for the generation of hepatocyte lumina

Given the strength of the phenotype and because Rab35 had no previous connection to hepatocyte lumen morphology, we explored its function in more detail. To begin with, we validated the specificity of the Rab35 RNAi phenotype. First, out of six designed siRNAs, five yielded Rab35 mRNA down-regulation above 50% after 96h and showed various degrees of lumen alteration (Extended data Fig.4d-e). The three siRNAs (siRab35 #2, #4, #5) that consistently yielded the strongest phenotype reduced Rab35 mRNA (Extended data Fig.4e) and protein level more than 70% (Figure 3c). Second, we rescued the Rab35 RNAi phenotype by expressing human Rab35, which is resistant to siRab35 #4. We quantified the effect of Rab35 knock-down on lumen morphology by measuring the radius of individual lumina in the control and knock-down conditions, and plotting the frequency distribution of values. There was a consistent shift towards larger lumina in the knock-down conditions by the three siRNAs targeting Rab35 mRNA (Figure 3d). Importantly, whereas in control conditions lumina barely had a radius larger than 6 μm, upon Rab35 silencing ^~^ 20–25% of lumina had a radius > 6 μm. Re-expression of human EGFP-Rab35 rescued the phenotype, shifting the distribution of lumen radius towards the control, whereas expression of EGFP had no affect (Figure 3e).

The change in cell polarity induced by depletion of Rab35, raises the question of whether it is required for formation of the apical bulkheads. Rab35 was enriched in the apical surface as well as lateral plasma membrane and cytoplasmic vesicles (Figure 3f), in line with its endosomal localization^32,33^. Upon silencing, this staining was markedly reduced (Figure 3f). Expression of exogenous EGFP-tagged Rab35 yielded a similar pattern of localization (Extended data Fig.4f). However, EGFP-Rab35 was not only enriched apically but also present on the transversal connections which were dynamically remodelled as the apical lumen expanded anisotropically.

### Loss of apical bulkheads and cyst formation upon Rab35 KD via cell self-organization

To gain insights into the change in polarity and lumen morphogenesis, we imaged LifeAct-EGFP expressing cells transfected with Rab35, or Luciferase siRNA as control, by live-cell time-lapse microscopy. Whereas normal and control differentiating hepatoblasts formed elongated lumina (e.g. Figure 1c), upon Rab35 depletion, they generated spherical lumina, initially between two cells (Figure 3g, Video 8). With time, we observed major cell rearrangements, whereby cells moved and reshaped their apical surface leading to the fusion of lumina and the formation of 3D multi-cellular cysts (Extended data Fig.5a, Video 9), similar to Figure 3b. Again, such a re-organization was not a result of cell division, as for other cysts formed *in vitro*^1^, but rather by a self-organization process. A spherical expansion of the lumen occurred only in the cases where the cells failed to form the striped actin-rich bulkheads-pattern indicative of the BC lumina (Figure 3g, Video 8 and 9). Conversely, the elongated lumina that still formed always contained the transversal actin stripes. Careful inspection of the live cell imaging videos (e.g. Video 1 and 9) indicated that the disappearance of the transversal bulkheads precedes the formation of a spherical lumen.

To corroborate the loss of the bulkheads in the spherical lumina induced by Rab35 KD, we examined their ultra-structure by EM on serial sections and 3D reconstruction of the entire lumen volume. We focused on large cyst-like lumina formed by several cells. Individual EM sections of a cyst-like lumen between five cells showed that the bulkheads that are normally present in the BC lumina were absent (Figure 3h, Extended data Fig.5b). This was confirmed by the 3D model of the lumen based on rendering plasma membranes and tight junctions (Figure 3i). In addition, the tight junctions between the cells did not protrude into the lumen, as seen at the sagittal cross section of the 3D model (Figure 3i).

### Re-engineering of liver tissue architecture by silencing of Rab35 *in vivo*

If the transversal bulkheads confer to hepatocytes their specific polarity and, consequently, the cell-level anisotropic growth of the apical lumen, one could exploit their loss to re-engineer liver tissue, i.e. to predictably modify its structure, particularly the geometric characteristics of the BC network. The structure of liver tissue depends on two types of cell polarity, the polarity of hepatocytes that leads them to form the BC and the simple polarity of cholangiocytes that form the bile ducts. Loss of the transversal bulkheads in hepatocytes *in vivo* should change cell polarity resulting in a re-organization of cell-cell interactions. If so, the BC should be replaced by bile duct-like simple epithelial tubes. The complete loss of Rab35 in a KO mouse line is embryonically lethal^34^, presumably due to cytokinesis defects^32^. To circumvent this problem and deplete Rab35 as *in vitro*, we took advantage of lipid nanoparticles (LNP) developed for human therapeutics, enabling the specific delivery of siRNAs to hepatocytes in the liver^35,36^. To target the E13.5 embryonic liver, we used a method for *in utero* injection via vitelline vein^37^. We first validated the technique on mice expressing membrane-targeted GFP. We performed the *in utero* injection of LNP-GFP or Luciferase (as control) siRNA in E13.5 embryos and collected the livers after 4 days of development (Extended data Fig.6a). The GFP signal in the liver was markedly and homogeneously reduced in hepatocytes, whereas different cell types, e.g. hematopoietic cells were unaffected (Extended data Fig.6b).

We next formulated the Rab35 siRNA validated *in vitro* (Figure 3c-f) and Luciferase siRNA into LNP, injected them into embryonic livers, and analysed the effect using a pipeline of immunostaining, deep tissue imaging and 3D reconstruction^8^ (Figure 4a). As in control liver, E17.5 livers injected with LNP-Luciferase siRNA developed normal elongated BC tubules formed by two adjacent hepatocytes (Figure 4a’). Strikingly, LNP-Rab35 siRNA-injection indeed induced the formation of large tubular structures in the liver parenchyma (Figure 4a’’). As these structures are remarkably similar to bile duct at this developmental stage, we needed to rule out that they may be formed by bile duct cells. First, the tubular structures are present throughout the parenchyma and distant from the portal area where the bile ducts are located. Second, staining for HNF4a (hepatocyte marker) and Sox9 (bile duct cell marker) confirmed that the cells forming the tubules expressed hepatocyte markers and were not fully differentiated bile duct cells (Figure 4a’’’, Extended data Fig.6c).

**Figure 4:**
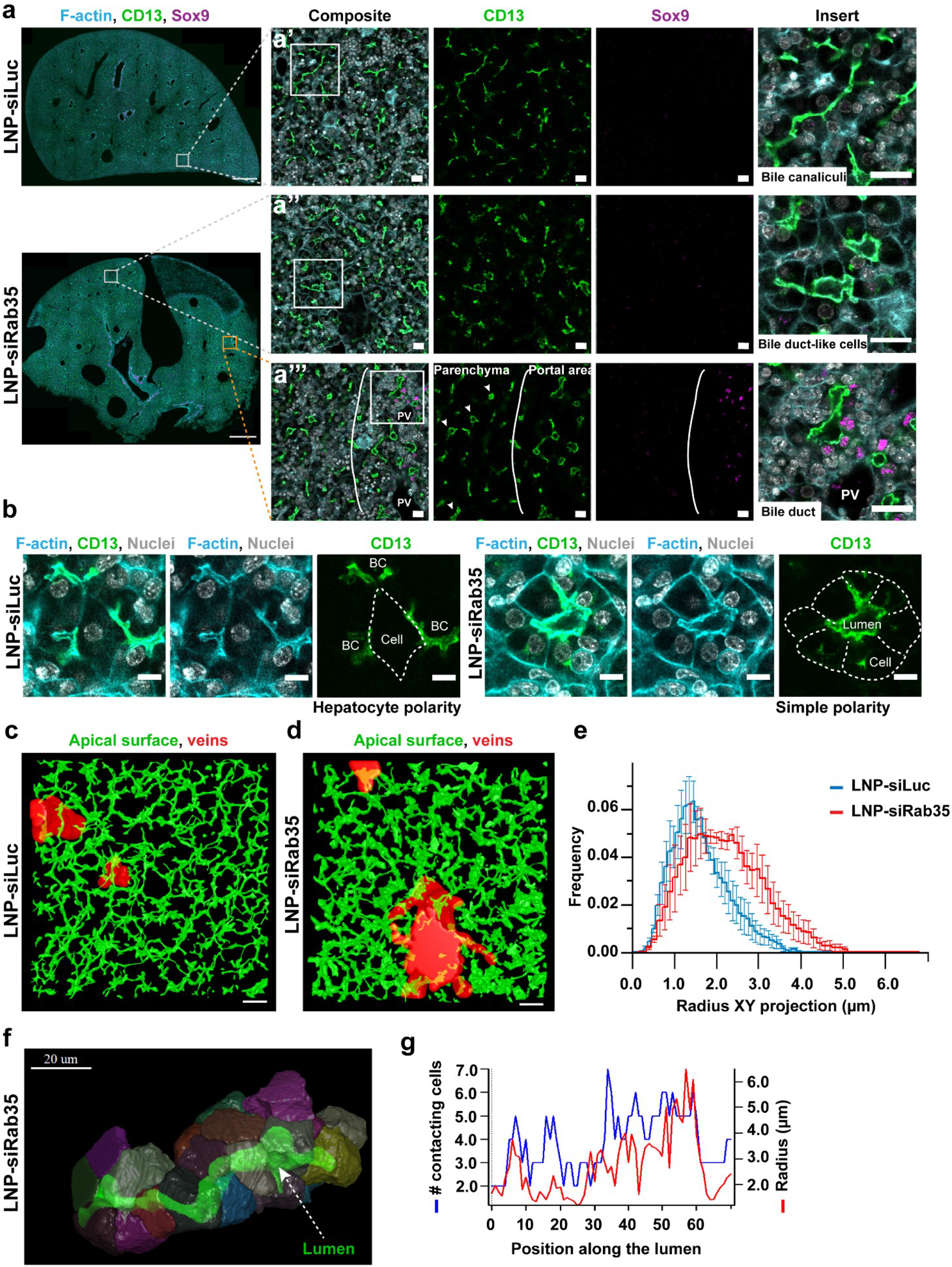
Silencing of Rab35 in vivo results in altered cell polarity and liver tissue architecture. **a**, Immunofluorescence images of liver tissue collected 4 days after in utero injection of Luciferase (siLuc) and Rab35 (siRab35) siRNAs formulated into LNP via vitelline vein in E13.5 embryos. The square on the low magnification images (scale bar: 500 μm) shows where the high resolution image was taken (scale bar: 20 μm). Imaged areas are located in the liver parenchyma, devoid of bile duct cell marker Sox9. The inserts (scale bar: 20 μm) in panels **a’** and **a’’** show the difference between BC and bile duct-like lumina in LNP-siRab35 injected liver. Panel **a’’’** compares tubular lumina in the parenchyma to the bile duct lumina in the portal area (Sox9-positive). **b**, Immunofluorescence images of liver tissues from **a** show examples of hepatocyte polarity in the control tissue (a single hepatocyte forms multiple lumina per cell) and simple apico-basal polarity in LNP-siRab35 injected liver (cell have a single apical domain oriented towards a shared lumen). **c**, **d**, 3D reconstruction of lumina labelled with an apical marker CD13 in 100 μm-thick sections of liver tissue injected with LNP-siLuc (**c**) and LNP-siRab35 (**d**). Scale bar: 30 μm. See also Video 10. **e**, Quantification of the lumen radius distribution based on the 3D reconstructions such as in **c** and **d** (n= 3, error bars: SEM). **f**, 3D reconstruction of a tubule found in LNP-siRab35 injected livers shows lumen in green surrounded by multiple cells. See also Video 11. **g**, Quantification of number of cells surrounding the lumen in relation to lumen radius and position along the tubule.

Importantly, the organization of the tubules suggested a re-arrangement of cell polarity. Whereas in control tissue individual hepatocytes formed multiple lumina per cell (characteristic of hepatocyte polarity), the cells forming the large tubular structures polarized with a single apico-basal axis and shared the same apical lumen (Figure 4b, see below).

3D reconstruction of apical surfaces (marked with CD13) in 100 μm-thick sections revealed the typical appearance of 3D BC network in normal and LNP-Luciferase siRNA-injected livers (Figure 4c, Video 10). In contrast, in LNP-Rab35 siRNA-injected livers, the 3D reconstruction showed profound changes in lumen morphology in proximity to the central vein, i.e. distant from peri-portal bile duct (Figure 4d, Video 10). The quantification from the reconstructed lumina showed a general increase in lumen radius (Figure 4e), similar to the one observed *in vitro* (Figure 3d). Remarkably, 30% more of BC lumina had radii larger than 2 μm compared to control livers (control 26.7% ± 9.4%, siRab35: 57% ± 11.9; mean ± SEM). 3D reconstruction of segments of large tubular structures revealed that the cells had a modified polarity, as the tubes had an average of 4 cells sharing the same lumen (Figure 4f-g; Video 11).

## Discussion

The molecular and physical mechanisms underlying the anisotropy of lumen formation are an emerging area of research. In this study, by searching for a mechanism that could explain the anisotropy of hepatocyte apical lumina, we discovered the existence of specific extensions of the apical membrane sealed by tight junctions in the lumen between two adjacent hepatocytes. The best analogy we could find for these structures are the bulkheads of boats, ships and planes. Bulkheads provide structural stability and rigidity, strengthening the structure of elongated vessels. From the physics of thin-shells, formation of a tubular lumen with inner pressure and no outlets, such as the forming BC, requires anisotropy of surface tension and/or rigidity of the wall^24,25^. The apical bulkheads are structural elements which can provide such anisotropy and mechanical stability to the elongating cylindrical lumen under inner pressure. Interestingly, the bulkheads follow a quasi-periodic pattern, whose distance is in the range of the diameter of the lumen, as in human-made constructions, where bulkheads are load-bearing structures. Here, they provide forces required for maintaining a non-spherical lumen. One can consider the cylinder with bulkheads as a “chain of spheres”, which is mechanically stable. Mechanistically, the position of the bulkheads could be determined by mechano-sensing mechanisms coupled to the tension and local curvature through the actin cortical mesh^38^. The elongation of the apical lumen entails also the movement and re-arrangement of cell-cell contacts which is accompanied by the formation of new bulkheads (Figure 1c, Extended data Fig.1). Upon loss of the bulkheads caused by Rab35 down-regulation the apical surfaces of hepatocytes lose their anisotropic growth and the elongated lumina convert into spherical.

We showed the apical bulkheads are present in embryonic liver, suggesting that they are not a cell culture artefact but have physiological relevance. In addition, the elongation assisted by apical bulkheads does not rely on cell division and therefore can explain the BC extension in quiescent differentiated hepatocytes in later stages of liver development^23^. Their dynamic and adaptable nature fit the requirements of a growing, branching and fusing BC network *in vivo*. Remarkably, we succeeded in re-engineering liver tissue structure by down-regulation of Rab35 *in vivo.* This resulted in the modification of the cell polarity of hepatocytes which, instead of forming BC, self-organized into tubular epithelial structures resembling bile ducts. It will be interesting to assess whether such morphological changes have consequences on hepatocyte cell fate and function.

We obtained several cues to the mechanisms underlying the apical bulkheads formation from the morphological analysis and functional screen by RNAi. First, the bulkheads are characterized by a T-shaped arrangement of tight junctions which seal the two halves of the bulkheads (Figure 1B, 2F). To our knowledge, this organization is unprecedented in polarized cells. Second, given that the tight junctions are connected to actin filaments, it is no surprise that the bulkheads contain F-actin transversally to the lumen elongation, thus introducing anisotropy in apical surface tension. Third, by a focused RNAi screen for established regulators of cell polarity we found that the small GTPase Rab35, a regulator of endosomal recycling^32,33,39^, is required for the formation of the apical bulkheads and hepatocyte lumen shape. Based on the previous work on Rab35 function^32,40^, we envision that it could regulate the intracellular distribution and function of apical recycling endosomes to deliver transmembrane proteins, e.g. junction components, at the site of bulkheads initiation and/or growth. The presence of clusters of vesicles at the base of the bulkheads as visualized by EM supports this view. In addition, Rab35 is also known to coordinate membrane trafficking with the organization of the actin cytoskeleton^40,41^ and its precise role in bulkhead formation awaits further research. Rab35 could regulate the nucleation and/or dynamics of the F-actin at the bulkheads, e.g. via known (e.g. MICAL1 and OCRL)^42–44^ or unknown hepatocyte-specific effectors. Also, the function of genes implicated in hepatocyte polarity, e.g. *Par1b*, *Pard3*, *Cldn2*, *Cldn3* and *Lkb1* (cAMP-Epac-MEK-AMPK pathway regulating BC network formation)^21,29,45–50^, should be re-visited specifically in the context of the bulkheads and anisotropy of lumen elongation.

Our data thus suggest that transversal mechanical coupling between hepatocyte apical surfaces underlies the formation of BC and provide new insights into the longstanding problem of lumen morphogenesis in embryonic liver.

## Acknowledgements

We are grateful to Arnaud Echard for discussions and sharing various reagents. We thank Meritxell Huch, Elisabeth Knust, Kai Simons, Ivan Baines, and Janelle Lauer for stimulating discussions and critical reading of the manuscript. We acknowledge Sandra Segeletz for sharing expertise in recombinant adenovirus production and live-cell imaging, and Alexandra Kalaidzidou for visualization artwork. We acknowledge the DRESDEN-concept Genome Center for the bulk RNA-sequencing service, the Core Facility of the CMCB Technology Platform at TU Dresden for the support of the Light Microscopy Facility, and the Center for Information Services and High Performance Computing (ZIH) at TU Dresden for generous allocation of computer time. We would like to thank the following Services and Facilities of the Max Planck Institute of Molecular Cell Biology and Genetics for their support: Antibody Facility, Biomedical Services, Electron Microscopy Facility, Light Microscopy Facility, Protein Expression, Purification and Characterization (PEPC) Facility, and Scientific Computing Facility, particularly Lena Hersemann and Noreen Walker.

This research was financially supported by the German Federal Ministry of Research and Education (BMBF) (LiSyM, grant number 031L0038), the European Research Council (ERC) (grant number 695646), eutsche Forschungsgemeinschaft (DFG, erman esearch Foundation) under Germany’s Excellence Strategy – EXC-2068– 390729961– Cluster of Excellence Physicsof Life of TU Dresden and the Max Planck Society (MPG).

## Author contributions

Conceptualization, M.Z., L.B. and Y.K.; Methodology, L.B., J.D. and S.S.; Investigation, L.B., U.R., J.D., S.S., J.I.V., C.F. and H.R.; Software, E.G., H.M-N. and J.I.V; Formal analysis, L.B., U.R., E.G., H.M-N., J.I.V, C.F. and Y.K.; Resources, T.Z., E.S., T.P., V.K. and M.V.; Visualization, L.B., U.R., E.G., M.V., J.I.V., H.M-N., C.F. and Y.K.; Validation, L.B.; Writing – Original draft, L.B.; Writing – Review and Editing, M.Z. and Y.K.; Funding acquisition, M.Z.

## Methods

### Lead Contact

Further information and requests for resources and reagents should be directed to and will be fulfilled by the Lead Contact, Marino Zerial.

### Materials Availability

All unique/stable reagents generated in this study are available from the Lead Contact with a completed Materials Transfer Agreement.

### Data and Code Availability

Bulk RNA-sequencing data will be deposited to the NCBI Gene Expression Omnibus (GEO) under accession number GEO upon acceptance. Code for the scripts will be made available upon acceptance.

#### Experimental Model and Subject Details

##### Animals and animal handling

Animal experiments were conducted in accordance with German animal welfare legislation in pathogen-free conditions in the animal facility of the MPI-CBG, Dresden, Germany. Mice were maintained in a conventional barrier animal facility with a climate-controlled environment on a 12-h light/12-h dark cycle, fed ad libitum with regular rodent chow. Protocols were approved by the Institutional Animal Welfare Officer (Tierschutzbeauftragter), and necessary licenses were obtained from the regional Ethical Commission for Animal Experimentation of Dresden, Germany (Tierversuchskommission, Landesdirektion Dresden). For primary hepatoblast isolations, embryonic livers were collected from time-pregnant (E13.5-E14.5) wild-type mice C57BL/6JOlaHsd (Harlan laboratories/Envigo, USA) or C57BL6/JRj (Janvier Labs, France), or transgenic lines LifeAct-EGFP^51^, ROSAmT/mG^52^ or the incross of the two transgenic lines. For *in utero* LNP injection experiments, the GFP-expressing embryos were generated by crossing of ROSAmT/mG female with PGKCre(J) male^53^. The transgenic or wild-type embryos were injected *in utero* via vitelline vein at E13.5 and livers collected at E16.5 – E17.5.

##### Primary hepatocytes

Primary hepatocytes were isolated from male 8-12 week-old mice according to the established protocol^54^ and processed immediately for RNA isolation.

##### QBI-239A

Cell line was cultured in DMEM High Glucose (Cat. No. 41966-029, Gibco) with 5% FBS (heat inactivated) and used for the production of adenovirus.

#### Method Details

##### Dlk1+ hepatoblast isolation

Hepatoblasts were isolated as Dlk1+ fraction using magnetic cell separation as reported^26^ with several changes. Timed-pregnant mice (E13.5-14.5) were sacrificed by cervical dislocation. 16-24 embryonic livers were collected, fragmented and incubated in Liver perfusion media (Thermo Fisher Scientific, Cat. No. 17701-038) for 20 min in a 37°C water bath. The liver pieces were digested in Liver Digest Medium (Thermo Fisher Scientific, Cat. No. 17703-034,) supplemented with 10 μg/ml DNAse I (cat. No. DN25, Sigma-Aldrich) for further 20 min. Erythrocytes were lysed in Red blood cell lysis buffer (155 mM NH_4_Cl, 10 mM KHCO_3_, 0.1 mM Na_4_EDTA, pH 7.4). Digested cells were incubated with blocking antibody Rat Anti-Mouse CD16/CD32 (BD Biosciences, Cat. No. 553142, 1:100) for 10 min, then with Anti-Dlk mAb-FITC (MBL, Cat. No. D187-4, 1:40) for further 15 min. After washing with a buffer (0.5% BSA, 2 mM EDTA in PBS), cells were incubated with Anti-FITC MicroBeads (Miltenyi Biotec, Cat. No. 130-048-701, 1:10) for 15 min and separated on a magnetic column (Miltenyi Biotec, Cat. No. 130-024-201) according to the manufacturer’s protocol.

##### Hepatoblasts culture and differentiation

Culture wells were pre-coated with Matrigel (BD Biosciences, Cat. No. 356231, 10% Matrigel in PBS) for 30 min at 37°C and washed with PBS. Dlk1+ enriched cells were seeded in Expansion media (DMEM/F-12, GlutaMAX™ supplement (ThermoFisher, Cat. No. 31331028), 10% FBS, 1x ITS-X (Gibco, Cat. No. 51500-056), 0.1 μM Dexamethasone (Sigma-Aldrich, Cat. No D1756-25MG,), 10 mM Nicotinamide (Sigma-Aldrich, Cat. No. N0636-100G), 10 ng/ml human HGF (in-house production), 10 ng/ml mouse EGF (in-house production). In 96-well plates, cells were seeded at the density 13 000 cells/well, in 24-well plates at the density 60 000 cells/well. 24h later, the cells were overlaid with Differentiation media (MCDB131, no glutamine, (Gibco, Cat. No. 10372019), 5% FBS, 2 mM L-glutamine (ThermoFisher, Cat. No. M11-004), 1x ITS-X (Gibco, Cat. No. 51500-056), 0.1 μM Dexamethasone (Sigma-Aldrich, Cat. No D1756-25MG)) containing Matrigel to the final 5%. Cells were cultured for 5 days at 37°C, 5% CO_2_ with one additional Differentiation media change. Dlk1+ cells from E14.5 livers contained ^~^ 10% cells positive for bile duct cell marker Sox9 and were used in the experiments to optimize the growth of bile duct cysts. For other experiments, Dlk1+ cells from E13.5 livers were used, as all the cells gave rise to hepatocytes with BC in the above culture conditions.

##### Live-cell time-lapse microscopy

For the live-cell video microscopy, LifeAct-EGFP^51^ and ROSAmT/mG^52^ mouse strains were crossed and EGFP+ embryos were collected for the Dlk1+ cells’ isolation. The lk1+ cells were plated (transfected with siRNA) and imaged from the day 3 of the culture at an epifluorescent microscope Zeiss Axiovert 200 M with an incubator (37°C, 5% CO_2_) using an 20x objective (NA 0.5) in 10 min intervals for approximately 52 h. To image the localization of EGFP-Rab35, the Dlk1+ cells were isolated from ROSAmT/mG embryos and transduced with a recombinant adenovirus (AdenoEGFP-Rab35) at day 2 of the culture. The cells were imaged on the day 3 in 5 min intervals for up to 24h.

##### Immunofluorescence staining and confocal imaging

Cultured cells were fixed with 3% PFA for 15 min at RT, washed 3x with PBS, permeabilized with 0.1% Triton X-100 in PBS for 5 min at RT, and blocked with 0.5% FBS in PBS for min 30 min at RT. Primary antibodies were diluted in the blocking solution, rat monoclonal anti-CD13 (Novus, Cat. No. NB100-64843, 1:500), rabbit polyclonal anti-ZO-1 (Thermo Fisher Scientific, Cat. NO. 40-2200, 1:200), rat monoclonal anti-Cytokeratin 19 (Sigma-Aldrich, Cat. No. MABT913, 1:500) and goat polyclonal anti-Albumin (Novus, Cat. No. NB600-41532, 1:200) and were incubated 1h at RT or overnight at 4°C. Secondary antibodies (and/or Phalloidin-Alexa dyes (Thermo Fisher Scientific, 1:250) and DAPI (1 mg/ml, 1:1000)) were incubated for 1h at RT. For staining with rabbit polyclonal anti-Rab35 (Antibody Facility MPI-CBG Dresden, H26952, 1: 1000), the cells were permeabilized with 0.05% saponin and blocked with 3% BSA in PBS instead. Finally, cells were washed with PBS and imaged on laser scanning confocal microscopes Olympus Fluoview 1000 (objectives 40x/0.9/air, 60x/1.2/water), Zeiss LSM 700 (objectives 40x/1.2/water, 20x/0.8/air).

##### Single-molecule localization microscopy

Single-molecule localization microscopy (SMLM) experiments were performed on a Nikon Eclipse Ti microscope, which is specified elsewhere in detail^55^. Prior to acquisition, samples were irradiated in epifluorescence illumination mode to turn emitters, which were out-of-focus in the acquisition HILO illumination scheme, into the dark state. The length of the acquisition was set to capture the majority of emitters, i.e. imaging was concluded when only a very minor number of active emitters was detectable. When a critically low spot density was first reached, an acquisition scheme of 1 frame with low 405 nm excitation (activation) followed by 5 consecutive frames with 641 nm excitation, was used. Typical acquisition lengths were 60000-200000 frames with 20 ms integration time and 641 nm excitation. Raw image stacks were analysed with rapidSTORM 3.2^56^. The FWHM was set as a free fit parameter, but in the limits of 275–650 nm, which corresponds to an axial range of approximately 1μm^57^, the fit window radius was set to 1200 nm, the intensity threshold to 1000 photons, while all other fit parameters were kept from the default settings in rapidSTORM 3.2. Linear lateral drift correction was applied by spatio-temporally aligning distinct structures to themselves. This was facilitated by color-coding of the temporal coordinate with the built-in tool.

##### Transmission electron microscopy

*In vitro* cultures of hepatoblasts grown in 24-well plates were fixed by adding warm 2% glutaraldehyde in 200 mM HEPES, pH 7.4 to the culture medium at a 1:1 ratio and incubated for 5 min at 37 °C. Then the fixative and medium mixture was replaced by adding fresh 1% glutaraldehyde in 200 mM HEPES, pH 7.4 and samples incubated at 37°C for another 2 hours, then at room temperature overnight. For resin embedding samples were post-fixed with 1% osmium tetroxide and 1.5% potassium ferricyanide for 1 hour on ice, then contrasted en-bloc with 2% aqueous uranyl acetate for 2 hours at room temperature, dehydrated with a graded ethanol series: 70-80-90-96%, each for 10 min, and 4x 100%, each for 15 min, progressively infiltrated with LX-112 epoxy resin (Ladd Research Industries) and eventually polymerized at 60°C for 2 days. The plastic of the plate was broken off to release resin disks with a cell monolayer on one side. Disks were cut into small pieces that were remounted for longitudinal sectioning.

To collect the mouse embryonic liver, a pregnant mouse was sacrificed and liver organs were dissected from embryos and cut into a few pieces, which were immersion fixed with 4% paraformaldehyde in 200 mM HEPES, pH 7.4, 1mM CaCl_2_ overnight. Before resin embedding, liver tissue was cut in small pieces and additionally fixed with 1% glutaraldehyde in 200 mM HEPES, pH 7.4. Tissue was processed as described above except that epon resin was used for embedding. Tissue was sectioned at random orientation.

Serial, 90-nm thin sections were cut using a Leica Ultracut UCT ultramicrotome and deposited on formvar-coated, slot, copper grids. Sections were contrasted with 0.4% lead citrate for 1 min and imaged in a Tecnai T12 transmission electron microscope (ThermoFisher), operated at 100 kV and equipped with an axial 2k CCD camera (TVIPS).

Z-stack of images of serial sections were aligned using a TrackEM2 plugin in Fiji^58^. The liver apical membrane, bile canaliculus lumen, and junctional complex were segmented on aligned image stacks using IMOD^59^ and Blender ^60^ in order to reconstruct a 3-D model.

##### siRNA design, synthesis, transfection

Design of siRNA was performed using in-house software, first by testing all available sequences on the specificity for the target in mouse transcriptome (RefSeq in Pubmed), followed by elimination of sequences with significant complementarity to mouse miRNA, GC content below 25% and higher than 75% and immune responsive ones (like UGU, UGUGU, etc.). In addition, sequences were filtered using Reynolds rules^61^. Six siRNAs with highest functionality score were selected (Table 2) and synthesized by solid phase phosphoramidite method, purified by IE-HPLC and verified by LC-MS^62^. Pyrimidines in the sense strand and before A in antisense strand (UA, CA dinucleotides) were 2’-O-methylated (shown by lower case letters in the sequence) and both strands were 3’-modified with phosphorothioate dithymidylate (TsT) to enhance nuclease stability. Working stocks were prepared by diluting siRNAs to 10 μM in 10 mM Tris.HCl, pH 7.5. si NAs were transfected using transfection reagent Lipofectamine™ NAiMAX (Thermo Fisher Scientific, Cat. No. 13778075) according to the reverse transfection protocol provided by the manufacturer. The final concentration per well was 10 nM si NA and 0.1 v/v% Lipofectamine™ RNAiMAX. Control Luciferase and GFP siRNA were previously published: Control siRNA Luciferase^36^ (sense 5’-cuuAcGcuGAGuAcuucGAdTsdT-3’, antisense 5’-UCGAAGuACUcAGCGuAAGdTsdT-3’), GFP siRNA^63^ (sense 5’-ACAUGAAGCAGCACGACUUTT-3’, antisense 5’-AAGUCGUGCUGCUUCAUGUTT-3’).

##### Protein extraction and Western blotting

Cultured cells were lysed for 20 min in ice-cold SDS lysis buffer (20 mM Tris-HCl, pH 7.5, 150 mM NaCl, 1 mM EDTA, 1 mM EGTA, 1% SDS, 1% NP-40 (IGEPAL CA-630), freshly added 1/1000 CLAAAP (Chymostatin, Leupeptin, Antipain, Aprotinin, APMSF, Pepstatin), 1/100 Phosphatase Inhibitor Cocktail 2 and 3 (Sigma Aldrich). Per condition, five wells of a 96-well plate were pooled together into total of 125 μl of the SDS lysis buffer. The lysates were sonicated for 3 min and spun at 13000 x g for 10 min, 4⁰C. Protein concentration was measured with DC^™^ Protein Assay (Bio-Rad, Cat. No. 500-0116,). The samples were separated on 15% SDS-PAGE and transferred onto nitrocellulose membrane. Membranes were blocked and incubated with primary antibodies rabbit polyclonal anti-Rab35 (Antibody Facility MPI-CBG Dresden, F18256, 1:1000) and mouse monoclonal anti-γ-tubulin (Sigma-Aldrich, Cat. No. T6557, 1:2000) and secondary HPR-conjugated antibodies (1:10000) in 5 % dry-milk, 10 mM Tris-HCl pH 8.0, 200 mM NaCl, 0.1 % Tween20. The bound antibody was detected with ECL™ Western Blotting detection kit (GE Healthcare, Cat. No. RPN2209) on Hyperfilm ECL (Amersham GE Healthcare). The quantification of Western blots was done with Image J^64^, statistics were calculated and plots were generated in R^65^.

##### RNA isolation and RT-qPCR

Total RNA was isolated using RNeasy Mini Kit (Qiagen, Cat. No. 74104 50,) including the DNase I (Qiagen, cat. No. 79254) treatment step. Cells were lysed with RTL buffer supplemented with DTT. cDNA was synthesized using ProtoScript^®^II First Strand cDNA Synthesis Kit (NEB, Cat. No. E6560S), following the manufacturer’s protocol with the andom Primer Mix and the NA denaturation step. qPCR was performed on Roche LightCycler^®^ 96 in 10-μl reactions using FastStart Essential DNA Green Master (Roche, Cat. No. 06402712001). A housekeeping gene Rplp0 was used as an endogenous reference gene. qPCR primer Rplp0 (Forward: 5’- AGATTCGGGATATGCTGTTGGC -3’, everse: 5’- TCGGGTCCTAGACCAGTGTTC -3’), qPCR primer Rab35 (Forward: 5’- TGTCAACGTCAAGCGATGG -3’, everse: 5’- GGTCATCATTCTTATTGCCCACT -3’). Normalized relative gene expression value and % knock-down was calculated using ΔΔCq method^66^, statistics were calculated and plots were generated in R^65^.

##### Bulk RNA-sequencing

Following samples were collected in 4 biological replicates: E14.5 Dlk1+ hepatobalsts isolated and immediately processed for RNA isolation, *in vitro* differentiated hepatocytes from E14.5 Dlk1+ hepatoblasts differentiated for 4 days in Differentiation media with 4% Matrigel grown on fibronectin coating, mature hepatocytes isolated from adult male mice following published protocols^54^ and immediately processed for RNA isolation. The integrity of RNA was measured by Agilent 2100 Bioanalyzer. Preferentially, only samples with the RNA integrity number (RIN) > 9.0 were used. 1 μg mRNA was isolated from the total RNA by poly-dT enrichment using the NEBNext Poly(A) m NA Magnetic Isolation Module according to the manufacturer’s instructions. Final elution was done in 15ul 2x first strand cDNA synthesis buffer (NEBNext, NEB). After chemical fragmentation by incubating for 15 min at 94°C the sample was directly subjected to the workflow for strand specific RNA-Seq library preparation (NEBNext^®^ Ultra™ RNA Library Prep Kit for Illumina^®^). For ligation custom adaptors were used (Adaptor-Oligo 1: 5’- ACA CTC TTT CCC TAC ACG ACG CTC TTC CGA TCT-3’, Adaptor-Oligo 2: 5’- P-GAT CGG AAG AGC ACA CGT CTG AAC TCC AGT CAC-3’). After ligation, adapters were depleted by an XP bead purification (Beckman Coulter) adding bead in a ratio of 1:1. Indexing was done during the following PCR enrichment (15 cycles, 65 °C) using custom amplification primers carrying the index sequence indicated with ‘NNNNNN’. (Primer1: Oligo_Seq AAT AT AC C ACC ACC GAG ATC TAC ACT CTT TCC CTA CAC GAC GCT CTT CCG ATC T, primer2: GTG ACT GGA GTT CAG ACG TGT GCT CTT CCG ATC T, primer3: CAA GCA GAA GAC GGC ATA CGA GAT NNNNNN GTG ACT GGA GTT. After two more XP beads purifications (1:1) libraries were quantified using Qubit dsDNA HS Assay Kit (Invitrogen). For Illumina flowcell production, samples were equimolarly pooled and distributed on all lanes used for 75bp single read sequencing on Illumina HiSeq 2500 resulting in on average 30 Mio sequenced fragments per sample.

##### Recombinant adenovirus production and rescue experiments

Recombinant adenovirus to express EGFP-fused Rab35 (Human RAB35 cDNA, transcript variant 1 (NM_006861.7)) was produced using AdEasy^™^ Vector System (Qbiogene) developed by^67^. A linker GGGGSGGGGS was introduced between EGFP and RAB35. RAB35 fragment with the linker extension was amplified from the Addgene plasmid #47424, a gift from Peter McPherson^68^, and subcloned into pEGFP-C3 vector (Clontech) using ScaI and BamHI restriction sites. The EGFP-linker-RAB35 fragment was cloned into a transfer vector pShutle-CMV (AdEasy Vector System, Qbiogene) using SalI and HindIII restriction sites. The recombinant transfer vector was linearized by PmeI and transformed into electro-competent *E.coli* strain BJ5183-AD-1 (Stratagene, Cat. No. 200157-11) for *in vivo* recombination with pAdEasy vector. A positive clone was amplified in *E. coli* DH5α and linearized with PacI prior the transfection into the packaging cell line QBI-293A (Qbiogene HEK-293A cell derivative). Virus was amplified and purified via OptiPrep-gradient (Iodixanol 60 w/v% solution, Axis Shield, Cat. No.1114542). The control EGFP only virus was produced similarly.

E13.5 Dlk1+ hepatoblasts were seeded and transfected as described above with Luc siRNA or Rab35 siRNA #4. 72h later, the cells were infected with the recombinant adenovirus (EGFP or EGFP-Rab35) at dilution 1/1000 and 1/100, respectively. The cells were cultured for 2 more days, fixed, and stained with Phalloidin-Alexa 647 and DAPI. From the acquired images, the rescue of the lumen phenotype was quantified.

##### *In utero* siRNA-LNP injection

For the use *in vivo*, siRNA oligos were formulated into lipid nanoparticles (LNPs) with C12-200 lipoid as previously described^69^. siRNA-LNPs were delivered *in utero* into E13.5 embryonic livers via vitelline vein as described elsewhere^37^. We optimized the concentration of siRNA-LNPs to 5 mg/kg body weight and the length of the treatment to 4 days using siRNAs-LNPs targeting GFP mRNA^63^ in ROSAmG embryos (generated from the cross of ROSA mG/mT x PGKCre(J) lines). The weight of the embryos was estimated based on the published results^70^. Briefly, the pregnant mice were anesthetized in a narcosis box with isoflurane at 5% then placed on a heated stage attached to a narcosis mask flowing isoflurane at 2-3%. Analgesia was ensured by injecting 4 mg/kg of Metamizol right before surgery and maintained by adding 1.33 mg/mL of the same drug in the drinking water until sacrifice. The abdomen of the mice was shaved and then sterilized with ethanol; the eyes protected from desiccation using hydration cream. The uterus was exposed via vertical laparotomy. The embryos were then injected with 5μL of LNPs at 5mg/kg. The success of the injection was assessed by blood clearance from the targeted vessel. Embryos of the same mother were randomly assigned to be non-injected, injected with control siRNA or injected with the targeting siRNA. The injections were performed using pulled needles from manually labelled glass capillaries. After injections, embryos were placed back in the abdomen and the peritoneal cavity was closed by suturing. The epidermis was then closed with surgical clips. At the end of the surgery the mice were placed close to a heating lamp and monitored until complete awakening. The livers were collected at E17.5.

##### Liver tissue staining with optical clearing

Embryonic livers were fixed by PFA immersion (4 % PFA, 0.1 % Tween20, PBS) for 2 hours at RT and overnight at 4 °C. The PFA was neutralized by overnight incubation in 50 mM NH_4_Cl in PBS. The livers were later stored in PBS at 4 °C until processing. The livers were mounted in 4% low-melting agarose in PBS and cut into 100 μm – thick sections at a vibratome (Leica VT1200S). For deep tissue imaging, tissue sections were permeabilized with by 0.5% TritonX100 in PBS for 1 hour at RT. The primary antibodies rat monoclonal anti-CD13 (Novus, NB100-64843, 1:500) and rabbit monoclonal anti-Sox9 (Abcam, Cat. No. ab185966, 1:500) were diluted in Tx buffer (0.2% gelatin, 300 mM NaCl, 0.3% Triton X-100 in PBS) and incubated for 2 overnights at RT. After washing 5 x 15 min with 0.3% TritonX-100 in PBS, the sections were incubated with secondary antibodies donkey anti-rat 568 (BIOTIUM, Cat. No. 20092, 1:1000), donkey anti-rabbit 647 (Thermo Fisher Scientific, Cat. No. A31573, 1:1000) and DAPI (1 mg/ml, 1:1000) and Phalloidin-Alexa488 (Thermo Fischer Scientific, Cat. No. A12379, 1:150) for another 2 overnights. After washing 5 x 15 min with 0.3% TritonX-100 in PBS and 3 x 1 min with PBS, the optical clearing started by incubating the slices in 25% fructose for 4 hours, continued in 50% fructose for 4 hours, 75% fructose overnight, 100% fructose (100% wt/v fructose, 0.5% 1-thioglycerol, 0.1M phosphate buffer pH 7.5) for 6 hours, and finally overnight in SeeDB solution (Ke et al., 2013) (80.2% wt/wt fructose, 0.5% 1-thioglycerol, 0.1M phosphate buffer). The samples were mounted in SeeDB.

#### Quantification and Statistical Analysis

##### 3D reconstruction of bile canaliculi

Optically cleared 100-μm liver sections were imaged with an upright multiphoton laser-scanning microscope (Zeiss LSM 780 NLO) equipped with Gallium arsenide phosphide (GaAsp) detectors. Liver slices were imaged twice at low (20x/0.8 Zeiss objective) and high resolution (63x/1.3 Zeiss objective, 0.3 μm voxel size), respectively. Low-resolution overviews of the complete liver sections were created and used to select for regions where enlarged apical membranes were apparent. Selected regions (^~^300 μm x 300 μm x 100 μm; x, y, z) were then acquired at high resolution. High-resolution images were processed and bile canaliculi segmented, based on CD13 staining, with the Motion Tracking software as described (Morales-Navarrete et al., 2015; Morales-Navarrete et al., 2016). Local lumen radius distribution was calculated by assuming a maximal radius of 10 μm.

For cells segmentation, a selected region of an image (^~^70 μm x 70 μm x 60 μm; x, y, z) was denoised using the PURE-LET method^73^, i.e. through the ‘Pure enoise’ plugin in ImageJ, with Cycle-spin = 10 and Multiframe = 11. Shading and uneven illumination were then corrected using BaSiC algorithm^74^ and Rolling Ball Background Subtraction plugins in Fiji, respectively. The pre-processed image was imported to Motion Tracking and apical membranes were reconstructed as above. Cells surrounding an apical tube were segmented using the 3D active mesh approach with phalloidin staining as a marker of cells border, as described in (Morales-Navarrete et al., 2015).

##### Lumen radius quantification

To quantify the effect of Rab35 silencing and Rab35 rescue on lumen morphology *in vitro*, a custom script was written for FIJI to segment lumina on microscopy images based on the actin signal (Phalloidin-Alexa 647) and extract region statistics. For the rescue experiment, the segmentation mask was set so that only lumina with a minimum (70%) overlap with GFP channel (expressed protein) were kept for the analysis (the cells that actually express the protein). The script contained a pause for segmentation verification and manual correction. For quantifying lumen radius, we used “local thickness” as descriptor, which can be computed with a Fiji plugin (https://imagej.net/Local_Thickness). The local thickness at any interior point of an object is defined as the diameter of the largest circle that contains the point and completely fits into the object. For each lumen, the local thickness histogram, as well as the average local thickness, was computed. Then, the local thickness histogram of each object was normalized. To account for the different size of the objects, each normalized histogram was multiplied by a weighting factor *w_i_*, which is proportional to the estimated volume of the object i. Without losing generality, we defined 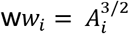, where *A_i_* is the number of pixels belonging to the object. Then, the histograms of all the objects in each image were summed up and normalized (i.e. to discard the effect of differences in the total amount of apical membrane between images). Finally, the averaged histograms (first over different images and then between different experiments N =3) are reported. Error bars show the standard error of the mean (SEM) per bin. The histogram quantification was performed using MATLAB R2020b.

##### Gene expression analysis

Basic quality control of raw sequencing data was performed with FastQC v0.11.2^75^. Reads were mapped to the mouse genome reference assembly GRCm38 and genes of the Ensembl release v92^76^ were quantified using STAR v2.5.2b^77^. The read duplication level was assessed using MarkDuplicates from Picard tools v2.10.2^78^, and dupRadar v1.8.0^79^. The count data of the samples were filtered for genes with more than 10 counts in any of the samples and served as input for DESeq2 v1.22.2^80^ to identify differentially expressed genes using a log2fold-change threshold of 1 and an adjusted p-value cut-off of 0.01. The heatmap was generated using R package gplots (function heatmap.2).

---

**Extended data Fig.1:**
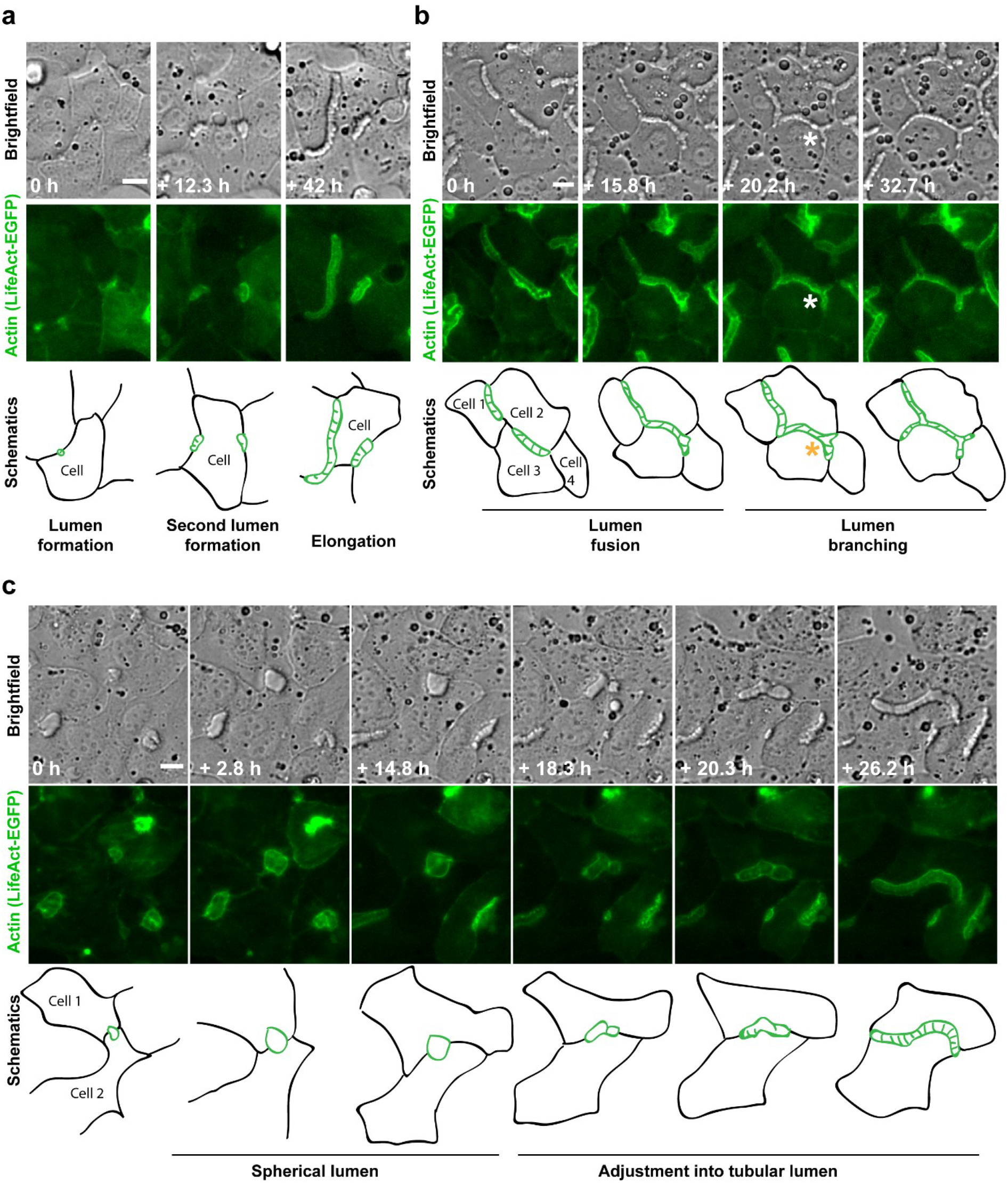
Live-cell video microscopy images of BC morphogenesis in LifeAct-EGFP expressing cells. **a**, A single polarizing differentiating hepatoblast forming multiple tubular lumina. Scale bar: 10 μm. See also Video 2. **b**, BC network forms between neighbouring cells by fusion and branching (star) of elongated tubular lumina. Scale bar: 10 μm. See also Video 3. **c**, A lumen formed between two cells starts growing spherical, but is later adjusted and continue to elongate as a typical BC lumen. Scale bar: 10 μm. See also Video 4.

**Extended data Fig.2:**
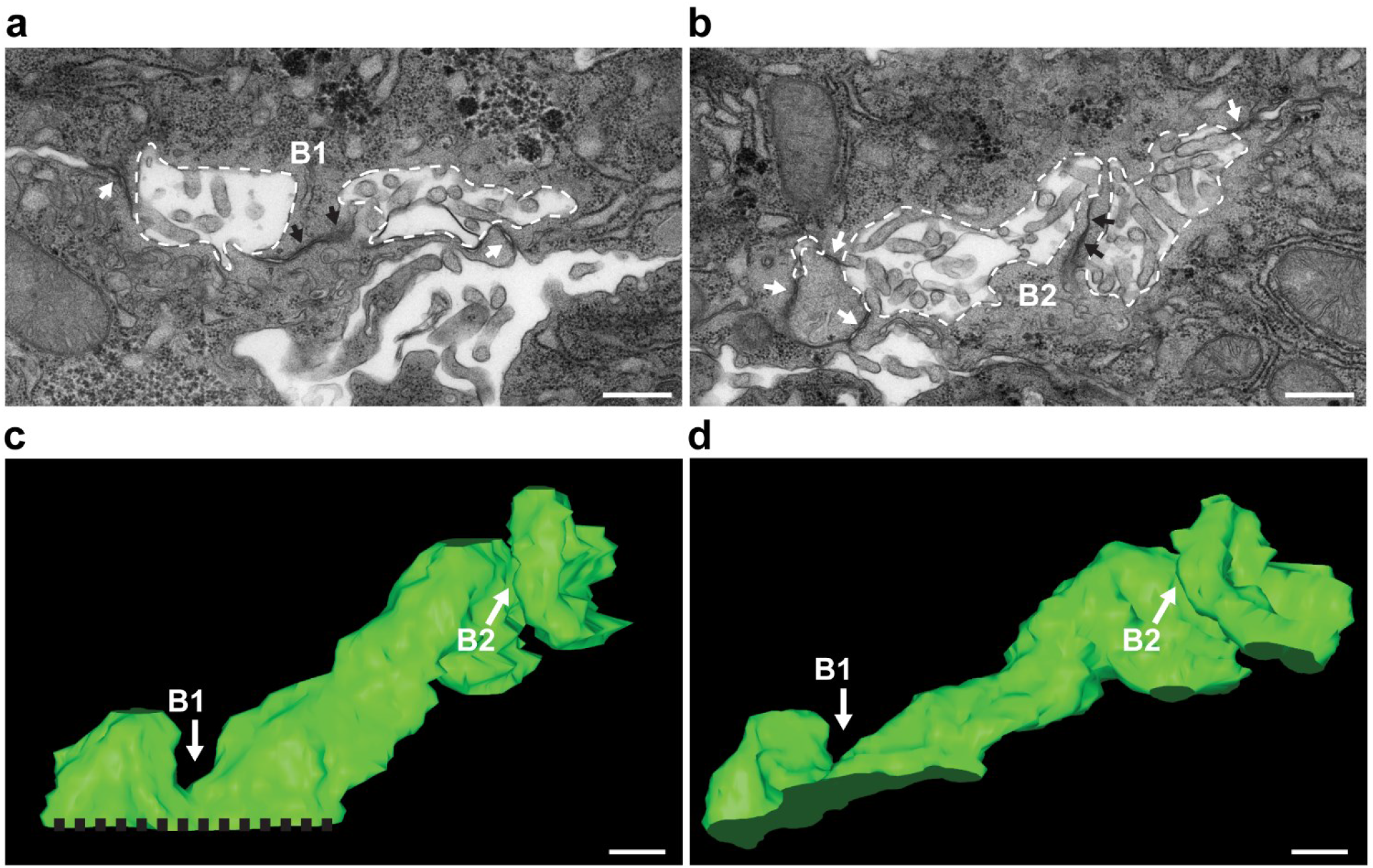
Transversal apical membrane structures form during BC lumen morphogenesis in embryonic liver. **a**, **b**, Electron microscopy images of two serial sections of a forming BC in a E15.5 liver, surrounded by tight junctions (white arrows). The lumen of the two adjacent hepatocytes is outlined in white. The black arrows point to the tight junctions within the bulkhead-like membrane connections B1 and B2. Scale bar: 500 nm. **c**, **d**, 3D model of the lumen in **a,** and **b,** based on lumen surface rendering on serial sections. At the left bottom side (dashed black line), the BC continues. White arrows indicate the bulkheads B1, B2 shown in **a** and **b** respectively. Scale bar: 500 nm.

**Extended data Fig.3:**
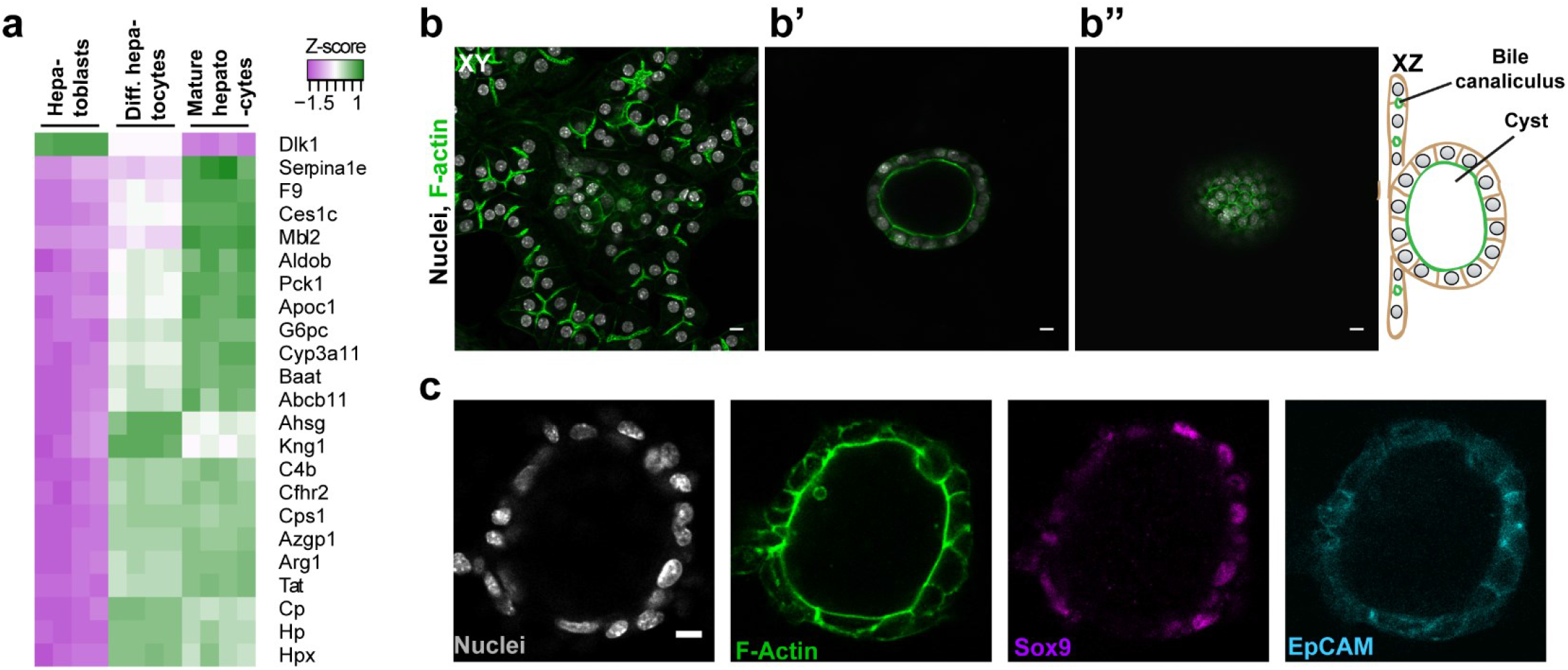
Validation of the culture system. **a,** *In vitro* differentiated hepatocytes express mature hepatocyte markers and down-regulate the hepatoblast marker Dlk1. Heatmap comparing the expression of selected hepatocyte marker genes in primary Dlk1+ hepatoblasts (Hepatoblasts), in vitro differentiated hepatocytes (Diff. hepatocytes) and control mature hepatocytes isolated from adult mouse livers (Mature hepatocytes). RNA-seq experiment in 4 biological replicates. **b**, A mixture of primary hepatoblasts and bile duct cells form BC and cysts under the same culture conditions. Images at different z-positions demonstrate that cysts grow in the z-direction (**b’**, **b’’**), while hepatocytes with BC form a cell-layer close to the well bottom. Cells stained for F-actin with Phalloidin-Alexa488. Scale bar: 10 μm. Schematics represent XZ view. **c**, Immunofluorescence staining for bile duct markers Sox9 and EpCAM in cyst-forming cells. Scale bar: 10 μm

**Extended data Fig.4:**
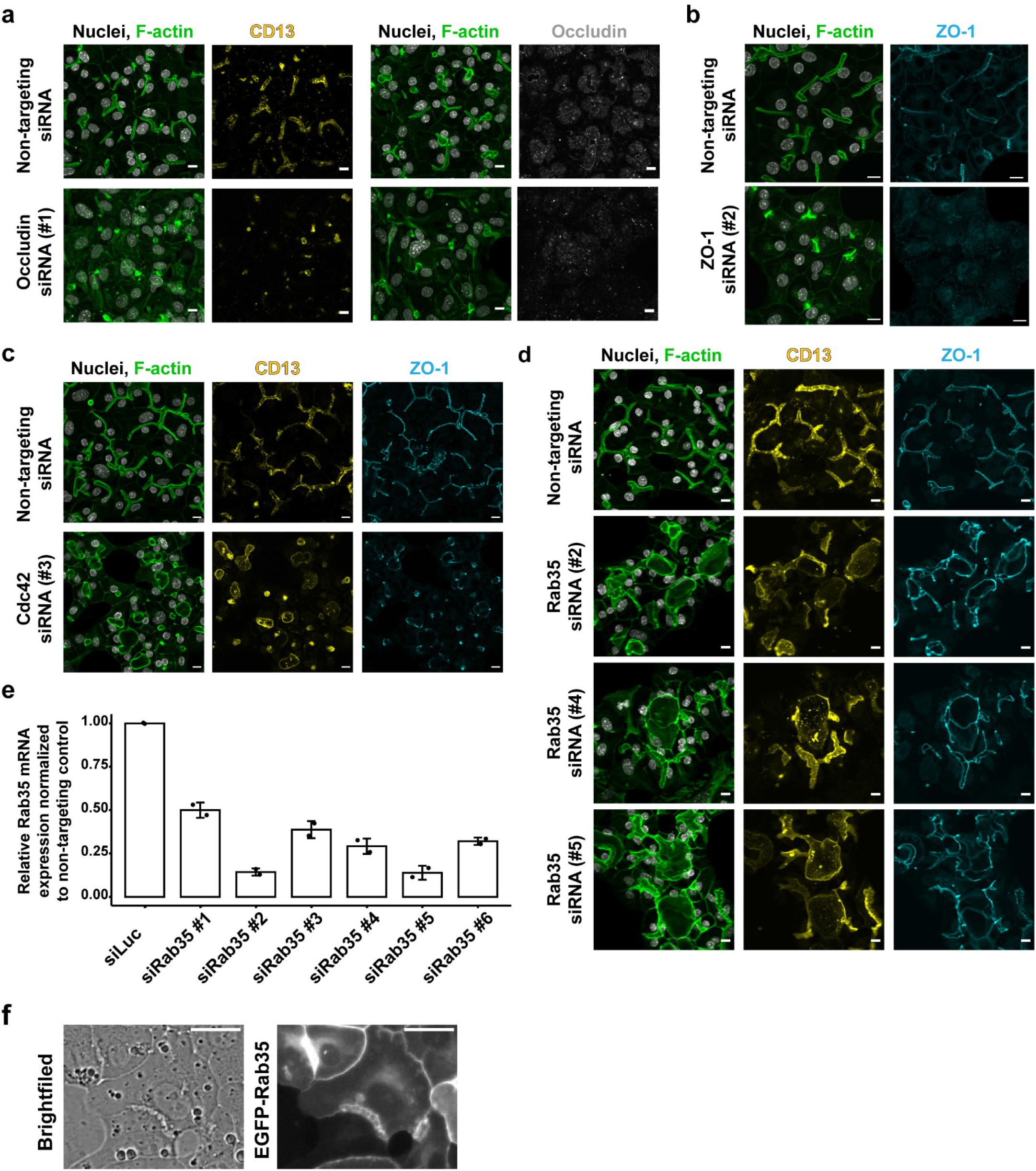
Conversion of hepatocyte polarity into simple apico-basal polarity. **a,** Down-regulation of occludin impairs the lumen formation in differentiating and polarizing hepatobalsts. Immunofluorescence images of cells stained for the apical marker CD13 and occludin. Scale bar: 10 μm. **b**, Down-regulation of the tight junction protein ZO-1 impairs the lumen formation in differentiating and polarizing hepatobalsts. Immunofluorescence images of cells stained for ZO-1. Scale bar: 10 μm. **c,** Down-regulation of Cdc42 leads to spherical lumina instead of BC in polarizing hepatoblasts. The polarity is not perturbed, as apical markers CD13 and ZO-1 still localized to the formed lumina. Scale bar: 10 μm. **d,** Down-regulation of Rab35 leads to profound changes in lumen morphology of polarizing and differentiating hepatoblasts. The cells form multicellular structures with a shared lumen positive for apical markers CD13 and ZO-1. Immunofluorescence images of cells treated with three different siRNAs. Scale bar: 10 μm. **e**, Estimation of knock-down efficiency of six siRNAs designed to target Rab35 mRNA 96h post-transfection in differentiating and polarizing hepatoblasts *in vitro* (n =2, SD). **f**, Localization of exogenous EGFP-Rab35 in polarizing hepatoblasts. Scale bar: 10 μm.

**Extended data Fig.5:**
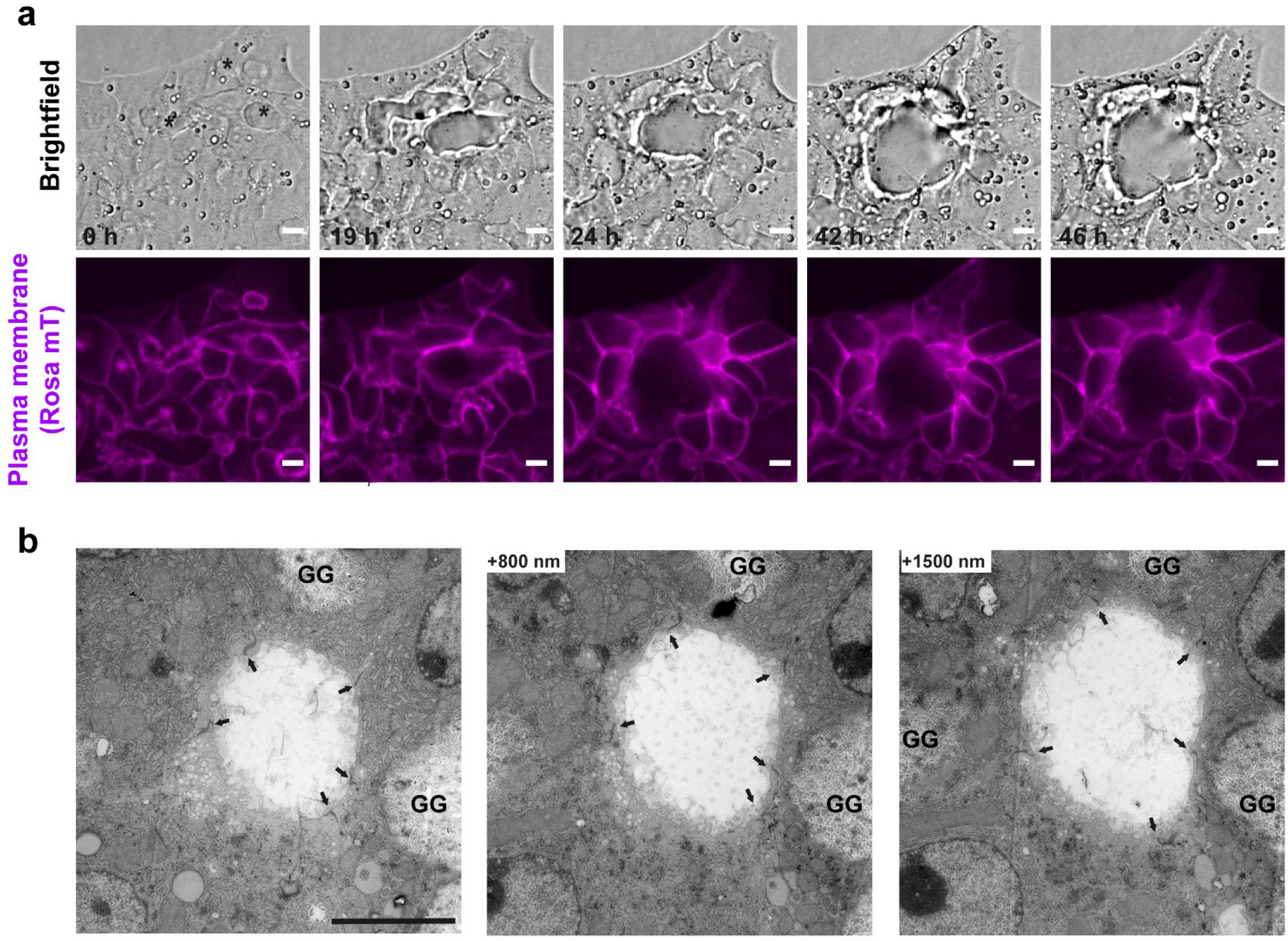
Silencing of Rab35 results in formation of spherical cysts via a cell self-organization process. **a**, Multicellular cyst-like structures form by cell re-arrangements. Images from the live-cell time-lapse microscopy experiment. The cells self-organize in such a way that the three separate lumina (black star) eventually fuse into one large spherical lumen, in the absence of cell division. Scale bar: 10 μm. See also Video 9. **b**, EM analysis of a cyst-like lumen resulting from the Rab35 knock-down. A series of longitudinal 90-nm sections of the lumen formed between five cells. The bulkheads typical for hepatocyte BC are absent in the lumen. Arrows indicate tight junctions. GG: glycogen granules. Scale bar: 5 μm.

**Extended data Fig.6:**
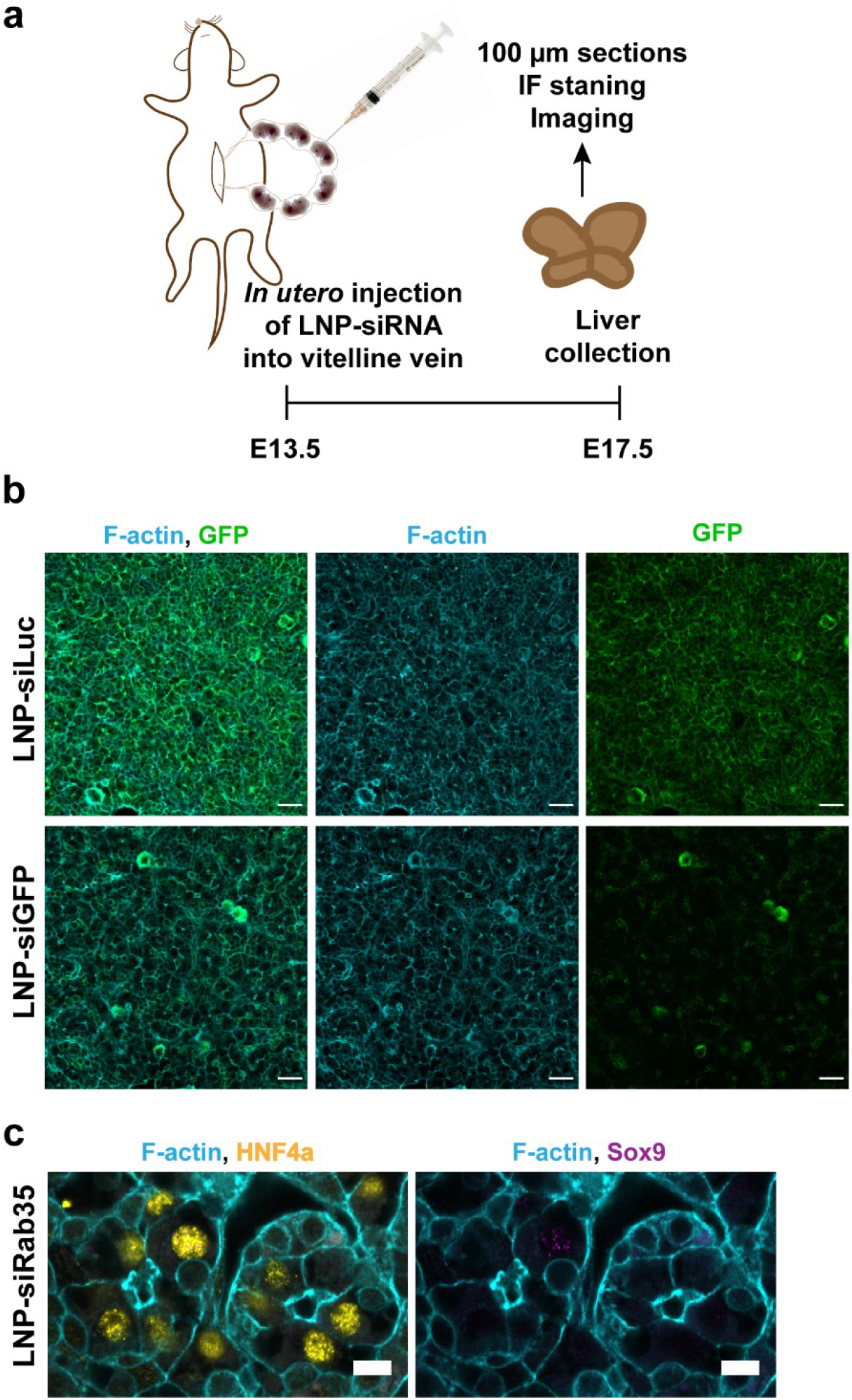
*In utero* injection method validated by silencing of GFP in GFP-expressing mouse line.

**a**, Schematic overview of *in utero* injection experiments. **b**, Microscopy images showing the down-regulation of GFP in GFP expressing mouse livers via *in utero* injection of LNP-siGFP in comparison to livers injected with control LNP-siLuc. Scale bar: 30 μm. **c**, Immunofluorescence microscopy images of the livers injected with LNP-siRab35 and stained for HNF4a (yellow) and Sox9 (magenta). Scale bar: 30 μm.

## Supplemental video and tables

**Video 1: Formation of BC *in vitro***

Live-cell time-lapse microscopy documenting the formation of BC between two differentiating hepatoblasts expressing LifeAct-EGFP. Images acquired in 10-min intervals. Scale bar: 10 μm.

**Video 2: Formation of multiple BC lumina by a single differentiating hepatoblast**

Live-cell time-lapse microscopy documenting the formation of multiple BC lumina by a single differentiating hepatoblasts expressing LifeAct-EGFP. Images acquired in 10-min intervals. Scale bar: 10 μm.

**Video 3: Branching and fusion of BC lumen**

Live-cell time-lapse microscopy of branching and fusing BC lumina formed by differentiating hepatoblasts expressing LifeAct-EGFP. Images acquired in 10-min intervals. Scale bar: 10 μm.

**Video 4: Recovery of tubular BC lumen**

Live-cell time-lapse microscopy documenting the adjustment of a spherical lumen into a tubular lumen in differentiating hepatoblasts expressing LifeAct-EGFP. Images acquired in 10-min intervals. Scale bar: 10 μm.

**Video 5: 3D model of BC based on EM data**

3D reconstruction of EM serial sections in Figure 2b based on apical plasma membranes and tight junctions rendering. The apical plasma membrane of the lumen-forming cells is in green and blue, the tight junctions are highlighted in red.

**Video 6: Animation of a simplified model of BC with bulkheads**

Simplified model of BC based on the 3D reconstruction in Figure 2e with periodic bulkhead-like membrane connections containing tight junctions (red) formed from the top or the bottom of the lumen. The apical plasma membrane of the lumen-forming cells is represented in green and blue.

**Video 7: Cysts formed upon Rab35 silencing *in vitro***

Confocal z-stack shows multiple cyst-like structures formed upon Rab35 silencing in differentiating hepatoblasts *in vitro*. Cells were stained for F-actin and nuclei. Scale bar: 30 μm.

**Video 8: Formation of a spherical lumen upon Rab35 silencing *in vitro***

Live-cell time-lapse microscopy showing the growth of a spherical lumen between two differentiating hepatoblasts expressing LifeAct-EGFP upon Rab35 silencing. Images acquired in 10-min intervals. Scale bar: 10 μm.

**Video 9: Formation of a multi-cellular cyst *in vitro* upon Rab35 silencing**

Live-cell time-lapse microscopy documenting the formation of a multicellular cyst upon Rab35 silencing in differentiating hepatoblasts expressing LifeAct-EGFP. Images acquired in 10-min intervals. Scale bar: 10 μm.

**Video 10: 3D reconstruction of luminal network in livers injected with LNP-siLuc or LNP-siRab35**

3D reconstruction of lumina labelled with an apical marker CD13 in 100 μm-thick sections of liver tissue injected with LNP-siLuc and LNP-siRab35. CD13 staining is shown first, then the 3D reconstruction based on the staining. The veins are shown in red. Scale bar: 30 μm.

**Video11: 3D reconstruction of a tubule formed in LNP-siRab35-injected liver**

3D reconstruction of a tubule found in LNP-siRab35 injected livers. First, CD13 staining is shown, next, the reconstructed lumen in green, and finally reconstructed cells in random colours. The lumen is surrounded by multiple cells.

**Table 2:siRNA sequences used in the study**

Excel spreadsheet

## References

1. Jewett, C. E. & Prekeris, R. Insane in the apical membrane: Trafficking events mediating apicobasal epithelial polarity during tube morphogenesis. Traffic 19, 666–678 (2018).

2. Navis, A. & Nelson, C. M. Pulling together: Tissue-generated forces that drive lumen morphogenesis. Seminars in Cell and Developmental Biology 55, 139–147 (2016).

3. Sigurbjörnsdóttir, S., Mathew, R. & Leptin, M. Molecular mechanisms of de novo lumen formation. Nature Reviews Molecular Cell Biology 15, 665–676 (2014).

4. Dasgupta, S., Gupta, K., Zhang, Y., Viasnoff, V. & Prost, J. Physics of lumen growth. Proc. Natl. Acad. Sci. U. S. A. (2018). doi:10.1073/pnas.1722154115

5. Stopka, A., Kokic, M. & Iber, D. Cell-based simulations of biased epithelial lung growth. Phys. Biol. 17, (2020).

6. Duclut, C., Sarkar, N., Prost, J. & Jülicher, F. Fluid pumping and active flexoelectricity can promote lumen nucleation in cell assemblies. Proc. Natl. Acad. Sci. U. S. A. 116, 19264–19273 (2019).

7. Treyer, A. & Müsch, A. Hepatocyte polarity. Compr. Physiol. 3, 243–87 (2013).

8. Morales-Navarrete, H. et al. A versatile pipeline for the multi-scale digital reconstruction and quantitative analysis of 3D tissue architecture. Elife 4, (2015).

9. Bryant, D. M. & Mostov, K. E. From cells to organs: Building polarized tissue. Nature Reviews Molecular Cell Biology 9, 887–901 (2008).

10. Andrew, D. J. & Ewald, A. J. Morphogenesis of epithelial tubes: Insights into tube formation, elongation, and elaboration. Developmental Biology 341, 34–55 (2010).

11. Datta, A., Bryant, D. M. & Mostov, K. E. Molecular regulation of lumen morphogenesis. Current Biology 21, (2011).

12. Müsch, A. The unique polarity phenotype of hepatocytes. Exp. Cell Res. 328, 276–83 (2014).

13. Müsch, A. From a common progenitor to distinct liver epithelial phenotypes. Curr. Opin. Cell Biol. 54, 18–23 (2018).

14. Ober, E. A. & Lemaigre, F. P. Development of the liver: Insights into organ and tissue morphogenesis. J. Hepatol. 68, 1049–1062 (2018).

15. Tanimizu, N. & Mitaka, T. Epithelial Morphogenesis during Liver Development. Cold Spring Harb. Perspect. Biol. 9, a027862 (2017).

16. Gissen, P. & Arias, I. M. Structural and functional hepatocyte polarity and liver disease. J. Hepatol. 63, 1023–37 (2015).

17. Tanimizu, N., Miyajima, A. & Mostov, K. E. Liver progenitor cells develop cholangiocyte-type epithelial polarity in three-dimensional culture. Mol. Biol. Cell 18, 1472–1479 (2007).

18. Prior, N. et al. Lgr5+ stem and progenitor cells reside at the apex of a heterogeneous embryonic hepatoblast pool. Dev. 146, (2019).

19. Antoniou, A. et al. Intrahepatic Bile Ducts Develop According to a New Mode of Tubulogenesis Regulated by the Transcription Factor SOX9. Gastroenterology 136, 2325–2333 (2009).

20. Tanimizu, N. et al. Intrahepatic bile ducts are developed through formation of homogeneous continuous luminal network and its dynamic rearrangement in mice. Hepatology 64, 175–188 (2016).

21. Wang, T., Yanger, K., Stanger, B. Z., Cassio, D. & Bi, E. Cytokinesis defines a spatial landmark for hepatocyte polarization and apical lumen formation. J. Cell Sci. 127, 2483–92 (2014).

22. Overeem, A. W., Bryant, D. M. & van IJzendoorn, S. C. D. Mechanisms of apical–basal axis orientation and epithelial lumen positioning. Trends Cell Biol. 25, 476–485 (2015).

23. Yang, L. et al. A single-cell transcriptomic analysis reveals precise pathways and regulatory mechanisms underlying hepatoblast differentiation. Hepatology 66, 1387–1401 (2017).

24. Landau, L. D. & Lifshitz, E. M. Theory of Elasticity, Third Edition: (Course of Theoretical Physics). Butterworth-Heinemann (1986).

25. Berthoumieux, H. et al. Active elastic thin shell theory for cellular deformations. New J. Phys. 16, (2014).

26. Tanimizu, N., Mitsuo Nishikawa, Hiroki Saito, Tohru Tsujimura, A. M. Isolation of hepatoblasts based on the expression of Dlk/Pref-1. J. Cell Sci. 116, 1775–1786 (2003).

27. Hannezo, E., Dong, B., Recho, P., Joanny, J. F. & Hayashi, S. Cortical instability drives periodic supracellular actin pattern formation in epithelial tubes. Proc. Natl. Acad. Sci. U. S. A. 112, 8620–8625 (2015).

28. Hayashi, S. & Dong, B. Shape and geometry control of the *Drosophila* tracheal tubule. Dev. Growth Differ. 59, 4–11 (2017).

29. Fu, D., Wakabayashi, Y., Ido, Y., Lippincott-Schwartz, J. & Arias, I. M. Regulation of bile canalicular network formation and maintenance by AMP-activated protein kinase and LKB1. J Cell Sci 123, 3294–3302 (2010).

30. Cohen, D., Brennwald, P. J., Rodriguez-Boulan, E. & Müsch, A. Mammalian PAR-1 determines epithelial lumen polarity by organizing the microtubule cytoskeleton. J. Cell Biol. 164, 717–27 (2004).

31. Yuan, H. et al. Hepatocyte-specific deletion of Cdc42 results in delayed liver regeneration after partial hepatectomy in mice. Hepatology 49, 240–249 (2009).

32. Kouranti, I., Sachse, M., Arouche, N., Goud, B. & Echard, A. Rab35 Regulates an Endocytic Recycling Pathway Essential for the Terminal Steps of Cytokinesis. Curr. Biol. 16, 1719–1725 (2006).

33. Klinkert, K., Rocancourt, M., Houdusse, A. & Echard, A. Rab35 GTPase couples cell division with initiation of epithelial apico-basal polarity and lumen opening. Nat. Commun. 7, 11166 (2016).

34. Dickinson, M. E. et al. High-throughput discovery of novel developmental phenotypes. Nature 537, 508–514 (2016).

35. Akinc, A. et al. Targeted delivery of RNAi therapeutics with endogenous and exogenous ligand-based mechanisms. Mol. Ther. 18, 1357–1364 (2010).

36. Zeigerer, A. et al. Rab5 is necessary for the biogenesis of the endolysosomal system in vivo. Nature 485, 465–470 (2012).

37. Ahn, N. J. et al. Intravenous and Intra-amniotic In Utero Transplantation in the Murine Model. J. Vis. Exp. (2018). doi:10.3791/58047

38. Meyer, K. et al. Bile canaliculi remodeling activates YAP via the actin cytoskeleton during liver regeneration. Mol. Syst. Biol. 16, (2020).

39. Mrozowska, P. S. & Fukuda, M. Regulation of podocalyxin trafficking by Rab small GTPases in 2D and 3D epithelial cell cultures. J. Cell Biol. 213, 355–69 (2016).

40. Klinkert, K. & Echard, A. Rab35 GTPase: A Central Regulator of Phosphoinositides and F-actin in Endocytic Recycling and Beyond. Traffic 17, 1063–1077 (2016).

41. Chua, C. E. L., Lim, Y. S. & Tang, B. L. Rab35 - A vesicular traffic-regulating small GTPase with actin modulating roles. FEBS Letters 584, 1–6 (2010).

42. Chaineau, M., Ioannou, M. S. & Mcpherson, P. S. Rab35: GEFs, GAPs and Effectors. Traffic 14, 1109–1117 (2013).

43. Dambournet, D. et al. Rab35 GTPase and OCRL phosphatase remodel lipids and F-actin for successful cytokinesis. Nat. Cell Biol. 13, 981–988 (2011).

44. Frémont, S. et al. Oxidation of F-actin controls the terminal steps of cytokinesis. Nat. Commun. 8, (2017).

45. Son, S. et al. Knockdown of tight junction protein claudin-2 prevents bile canalicular formation in WIF-B9 cells. Histochem. Cell Biol. 131, 411–424 (2009).

46. Grosse, B., Degrouard, J., Jaillard, D. & Cassio, D. Build them up and break them down. Tissue Barriers 1, e25210 (2013).

47. Slim, C. L. et al. Par1b Induces Asymmetric Inheritance of Plasma Membrane Domains via LGN-Dependent Mitotic Spindle Orientation in Proliferating Hepatocytes. PLoS Biol. 11, e1001739 (2013).

48. Homolya, L. et al. LKB1/AMPK and PKA Control ABCB11 Trafficking and Polarization in Hepatocytes. PLoS One 9, e91921 (2014).

49. Fu, D., Wakabayashi, Y., Lippincott-Schwartz, J. & Arias, I. M. Bile acid stimulates hepatocyte polarization through a cAMP-Epac-MEK-LKB1-AMPK pathway. Proc. Natl. Acad. Sci. U. S. A. 108, 1403–8 (2011).

50. Woods, A. et al. LKB1 is required for hepatic bile acid transport and canalicular membrane integrity in mice. Biochem. J. 434, 49–60 (2011).

51. Riedl, J. et al. Lifeact mice for studying F-actin dynamics. Nature Methods 7, 168–169 (2010).

52. Muzumdar, M. D., Tasic, B., Miyamichi, K., Li, N. & Luo, L. A global double-fluorescent cre reporter mouse. Genesis 45, 593–605 (2007).

53. Lallemand, Y., Luria, V., Haffner-Krausz, R. & Lonai, P. Maternally expressed PGK-Cre transgene as a tool for early and uniform activation of the Cre site-specific recombinase. Transgenic Res. 7, 105–12 (1998).

54. Klingmüller, U. et al. Primary mouse hepatocytes for systems biology approaches: A standardized in vitro system for modelling of signal transduction pathways. IEE Proc. Syst. Biol. 153, 433–447 (2006).

55. Franke, C. et al. Correlative single-molecule localization microscopy and electron tomography reveals endosome nanoscale domains. Traffic 20, 601–617 (2019).

56. Wolter, S. et al. RapidSTORM: Accurate, fast open-source software for localization microscopy. Nature Methods 9, 1040–1041 (2012).

57. Franke, C., Sauer, M. & De Linde, S. Van. Photometry unlocks 3D information from 2D localization microscopy data. Nat. Methods 14, 41–44 (2016).

58. Cardona, A. et al. TrakEM2 software for neural circuit reconstruction. PLoS One 7, (2012).

59. Kremer, J. R., Mastronarde, D. N. & McIntosh, J. R. Computer visualization of three-dimensional image data using IMOD. J. Struct. Biol. 116, 71–76 (1996).

60. Blender, F. Blender v2.5-a 3D modeling and rendering package. Blender Foundation (2010). Available at: http://www.blender.org.

61. Reynolds, A. et al. Rational siRNA design for RNA interference. Nat. Biotechnol. 22, 326–330 (2004).

62. Farzan, V. M. et al. Automated Solid-Phase Click Synthesis of Oligonucleotide Conjugates: From Small Molecules to Diverse *N* -Acetylgalactosamine Clusters. Bioconjug. Chem. 28, 2599–2607 (2017).

63. Gilleron, J. et al. Image-based analysis of lipid nanoparticle-mediated siRNA delivery, intracellular trafficking and endosomal escape. Nat. Biotechnol. 31, 638–646 (2013).

64. Miller, L. Analyzing western blots with ImageJ. 1–19 (2010). Available at: https://lukemiller.org/index.php/2010/11/analyzing-gels-and-western-blots-with-image-j/.

65. 3.5.1., R. D. C. T. A Language and Environment for Statistical Computing. R Foundation for Statistical Computing 2, https://www.R-project.org (2018).

66. Haimes, J. & Kelley, M. emonstration of a ΔΔ C q Calculation Method to Compute Relative Gene Expression from qPCR Data. GE Healthcare, Tech Note 1–4 (2010). Available at: http://dharmacon.gelifesciences.com/uploadedfiles/resources/delta-cq-solaris-technote.pdf. (Accessed: 1st May 2019)

67. He, T. C. et al. A simplified system for generating recombinant adenoviruses. in Proceedings of the National Academy of Sciences of the United States of America 95, 2509–2514 (1998).

68. Allaire, P. D. et al. The Connecdenn DENN domain: a GEF for Rab35 mediating cargo-specific exit from early endosomes. Mol. Cell 37, 370–82 (2010).

69. Love, K. T. et al. Lipid-like materials for low-dose, in vivo gene silencing. Proc. Natl. Acad. Sci. U. S. A. 107, 1864–9 (2010).

70. Kulandavelu, S. et al. Embryonic and neonatal phenotyping of genetically engineered mice. ILAR J. 47, 103–117 (2006).

71. Ke, M. T., Fujimoto, S. & Imai, T. SeeDB: A simple and morphology-preserving optical clearing agent for neuronal circuit reconstruction. Nat. Neurosci. 16, 1154–1161 (2013).

72. Morales-Navarrete, H., Nonaka, H., Segovia-Miranda, F., Zerial, M. & Kalaidzidis, Y. Automatic recognition and characterization of different non-parenchymal cells in liver tissue. in Proceedings - International Symposium on Biomedical Imaging 2016-June, 536–540 (2016).

73. Luisier, F., Vonesch, C., Blu, T. & Unser, M. Fast interscale wavelet denoising of Poisson-corrupted images. Signal Processing 90, 415–427 (2010).

74. Peng, T. et al. A BaSiC tool for background and shading correction of optical microscopy images. Nat. Commun. 8, (2017).

75. Simon Andrews. Babraham Bioinformatics - FastQC A Quality Control tool for High Throughput Sequence Data. Soil 5, 47–81 (2020).

76. Zerbino, D. R. et al. Ensembl 2018. Nucleic Acids Res. 46, D754–D761 (2018).

77. Dobin, A. et al. STAR: ultrafast universal RNA-seq aligner. Bioinformatics 29, 15–21 (2013).

78. Broad Institute. Picard Tools. Broad Institute, GitHub repository version 2.18.2 (2018). Available at: http://broadinstitute.github.io/picard/. (Accessed: 1st August 2018)

79. Sayols, S., Scherzinger, D. & Klein, H. dupRadar: a Bioconductor package for the assessment of PCR artifacts in RNA-Seq data. BMC Bioinformatics 17, 428 (2016).

80. Love, M. I., Huber, W. & Anders, S. Moderated estimation of fold change and dispersion for RNA-seq data with DESeq2. Genome Biol. 15, 550 (2014).

